# Age Differences in Diffusivity in the Locus Coeruleus and its Ascending Noradrenergic Tract

**DOI:** 10.1101/2021.11.23.469621

**Authors:** Shai Porat, Francesca Sibilia, Josephine Yoon, Yonggang Shi, Martin J. Dahl, Markus Werkle-Bergner, Sandra Düzel, Nils Bodammer, Ulman Lindenberger, Simone Kühn, Mara Mather

## Abstract

The noradrenergic locus coeruleus (LC) is a small brainstem nucleus that promotes arousal and attention. Recent studies have examined the microstructural properties of the LC using diffusion-weighted magnetic resonance imaging and found unexpected age-related differences in fractional anisotropy - a measure of white matter integrity. Here, we used three datasets (Berlin Aging Study-II, N = 301, the Leipzig Study for Mind-Body-Emotion Interactions, N = 220, and Stockholm Sleepy Brain, N = 49), to replicate published findings and expand them by investigating diffusivity in the LC’s ascending noradrenergic bundle. In younger adults, LC fractional anisotropy was significantly lower, compared to older adults. However, in the LC’s ascending noradrenergic bundle, we observed significantly higher fractional anisotropy in younger adults, relative to older adults. These findings indicate that diffusivity in the LC versus the ascending noradrenergic bundle are both susceptible to microstructural changes in aging that have opposing effects on fractional anisotropy.

**Highlights:**

- Fractional anisotropy in the locus coeruleus was lower in younger adults
- Fractional anisotropy in the noradrenergic bundle was higher in younger adults
- Sleep deprivation may affect diffusivity in younger adults more than older adults

## 1. Introduction

The locus coeruleus (LC) is the brain’s primary source for noradrenaline (F. S. Giorgi et al., 2020; Khanday et al., 2016; Lee et al., 2018), influencing arousal and attention throughout the day (Aston-Jones & Waterhouse, 2016; Dahl, Mather, Sander, et al., 2020; Mather, 2020; Mather & Harley, 2016; McGregor & Siegel, 2010; Sara, 2009). The LC also has widespread cortical projections that are susceptible to neurodegeneration (Aston-Jones & Waterhouse, 2016; Loizou, 1969; Loughlin et al., 1982; Morris, McCall, et al., 2020). Notably, the human LC is the primary site of early abnormal tau pathology (Braak & Del Trecidi, 2015; Liu et al., 2020; Mather & Harley, 2016) and until recently, *in vivo* microstructural properties of the LC were mostly unexplored (Edlow et al., 2016; Edlow et al., 2012; Langley et al., 2020).

Recently, Langley et al. (2020) examined the diffusive properties of the LC using diffusion-weighted MRI. They observed *higher* fractional anisotropy in the LC of older adults, compared with younger adults. Fractional anisotropy is widely used as a measure of structural integrity (higher fractional anisotropy typically indicates healthier axons) and has a strong inverse correlation with mean or radial diffusivity (Bhagat & Beaulieu, 2004; Kantarci et al., 2017; Kochunov et al., 2012). With aging, older adults typically display lower fractional anisotropy and higher mean diffusivity in white and gray matter compared with younger adults (Kantarci, 2014; D. A. Medina & M. Gaviria, 2008; Rose et al., 2008; Sullivan & Pfefferbaum, 2006; A. N. Voineskos et al., 2012). In addition, grey matter also typically shows lower fractional anisotropy and higher mean diffusivity in Alzheimer’s disease (Kantarci, 2014; Rose et al., 2008; Weston et al., 2015). Langley’s findings are the opposite of typical white matter age effects, though speculated to be associated with an age-related reduction in LC structure.

Given the surprising nature of the increased fractional anisotropy seen in older adults’ LC compared with younger adults’ LC, we were interested in testing whether these age differences replicate in larger samples. In addition, we were interested in whether the diffusivity within the LC might reflect a biomarker for transient states of inflammation or neuronal activity. Animal research suggests that inflammation restricts fluid flow (Yi et al., 2019). In addition, neuronal activity tends to swell neurons, reducing the interstitial fluid-filled spaces between them and decreasing diffusivity (Abe et al., 2017; Iwasa et al., 1980; Le Bihan et al., 2006; Nunes et al., 2021; Svoboda & Syková, 1991). Sleep deprivation affects both neuronal activity and inflammation and so we also examined whether a sleep deprivation manipulation would affect LC fractional anisotropy.

Unmyelinated neurons and numerous innervations to blood capillaries may expose the LC to toxins throughout aging (Bekar et al., 2012; Giorgi et al., 2020; Raichle et al., 1975). During the waking day, the LC has a high constant spiking rate which accumulates oxidative stress in the mitochondria of LC neurons (Weinshenker, 2018). In addition, excess noradrenaline not repackaged into synaptic vesicles promotes LC tau pathology (Kang et al., 2020). A lack of sleep, in particular REM sleep when the LC is least active (Takahashi et al., 2010), results in LC and amygdala hyperactivity (Goldstein & Walker, 2014; Khanday et al., 2016). Larger LC volume has been observed in patients with pathological anxiety (Morris, Tan, et al., 2020), which may be related to enhanced connectivity between the LC and amygdala (Giustino et al., 2020).

Using two large datasets (Berlin Aging Study-II, N = 301, (Delius et al., 2015), and the Leipzig Study for Mind-Body-Emotion Interactions, N = 220, (Babayan et al., 2019) of healthy young and older adults, we examined whether we could replicate LC fractional anisotropy findings as reported by Langley, et al., and compare them with the ascending noradrenergic bundle, which originates in the LC. To characterize diffusion properties within the ascending noradrenergic bundle, we relied on a probabilistic atlas of bilateral ascending noradrenergic fiber bundles originating in the LC and terminating in the transentorhinal cortex based on data from the Human Connectome Project (Sun et al., 2020; Tang et al., 2018). To test whether diffusivity in the LC is sensitive to short-term changes, we turned to a third dataset, the Stockholm SLEEPY Brain, final N = 49, which provides an opportunity to further investigate LC diffusivity under the context of sleep deprivation (Akerstedt, 2016).

## 2. Methods

Demographics and MRI sequence information across all three datasets can be found in Tables 1 through Table 4. The first dataset we examined is the Berlin Aging Study II (BASE-II) (Bertram et al., 2014; Delius et al., 2015) from timepoint two. BASE-II information can be found online (https://www.base2.mpg.de/en). In short, BASE-II is a multi-disciplinary and multi-institutional longitudinal study sampling from Berlin’s population. Because the BASE-II study included LC-MRI contrast measures, we asked whether these measures were associated with measures of LC and noradrenergic bundle diffusivity. The LC-MRI index potentially reflects neuromelanin accumulation as a byproduct of NE synthesis. Hence, it is supposed to indicate functional NE-density within the LC. If a lower LC-MRI contrast indeed reflects impaired functionality of the LC-NE system, detrimental downstream effects on pathways connecting the LC to the entorhinal cortex might be expected. Hence, we expect lower LC-MRI contrast ratios to be associated with lower diffusivity. BASE-II LC-MRI contrast values were previously quantified in a different study (Dahl et al., 2019).

**Table 1.**
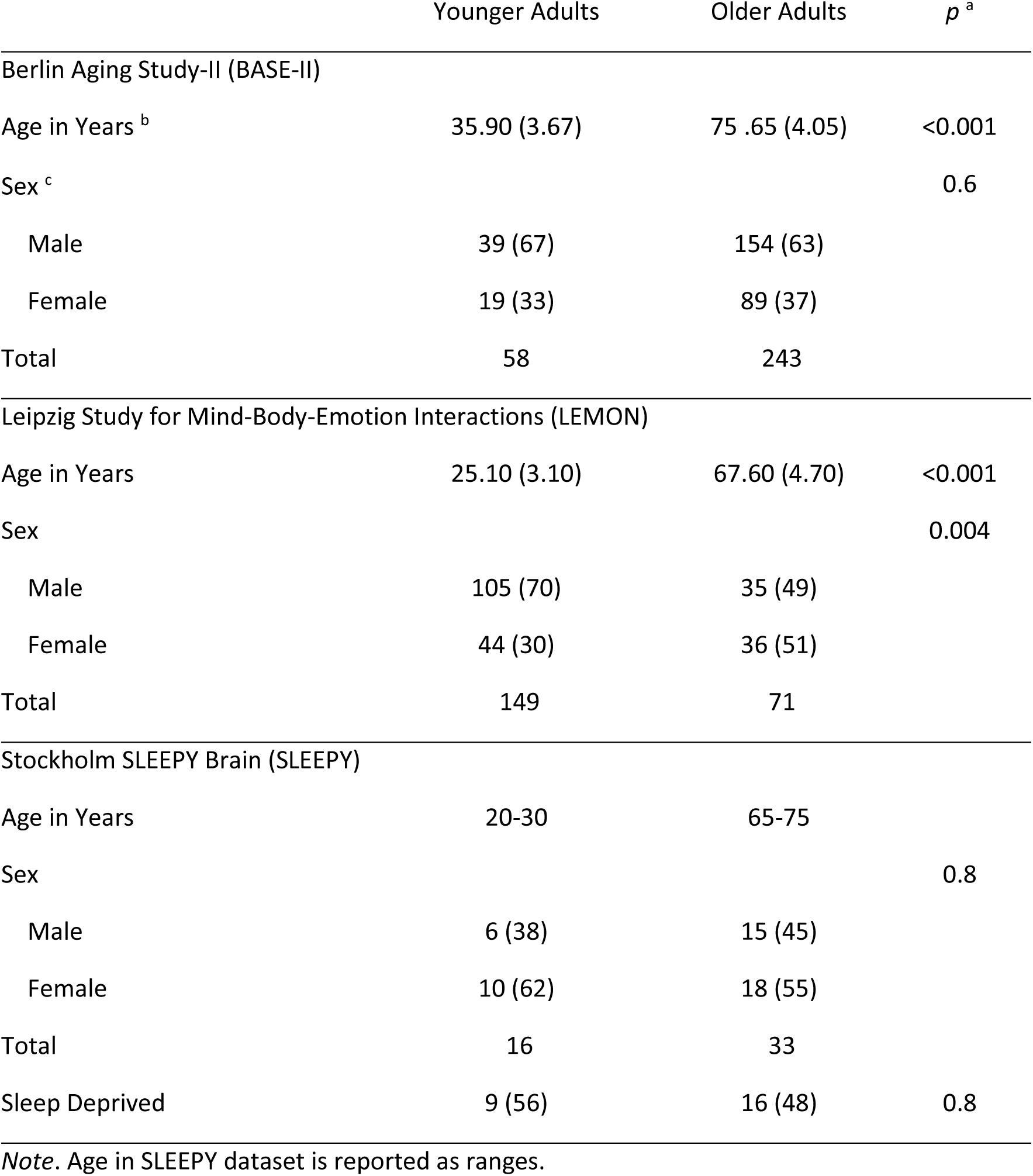
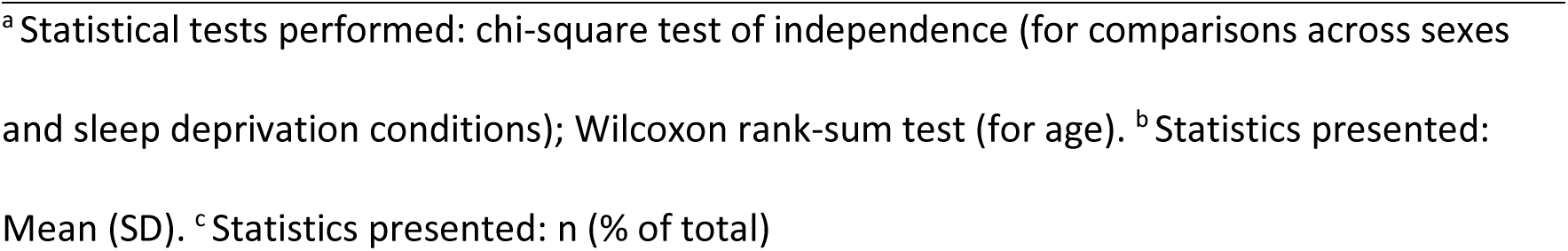
Demographics for Each Dataset

| | Younger Adults | Older Adults | $p^a$ |
| --- | --- | --- | --- |
| Berlin Aging Study-II (BASE-II) |  |  |  |
| Age in Years<sup>b</sup> | 35.90 (3.67) | 75.65 (4.05) | <0.001 |
| Sex<sup>c</sup> |  |  | 0.6 |
| Male | 39 (67) | 154 (63) |  |
| Female | 19 (33) | 89 (37) |  |
| Total | 58 | 243 |  |
| Leipzig Study for Mind-Body-Emotion Interactions (LEMON) |  |  |  |
| Age in Years | 25.10 (3.10) | 67.60 (4.70) | <0.001 |
| Sex |  |  | 0.004 |
| Male | 105 (70) | 35 (49) |  |
| Female | 44 (30) | 36 (51) |  |
| Total | 149 | 71 |  |
| Stockholm SLEEPY Brain (SLEEPY) |  |  |  |
| Age in Years | 20-30 | 65-75 |  |
| Sex |  |  | 0.8 |
| Male | 6 (38) | 15 (45) |  |
| Female | 10 (62) | 18 (55) |  |
| Total | 16 | 33 |  |
| Sleep Deprived | 9 (56) | 16 (48) | 0.8 |

*Note.* Age in SLEEPY dataset is reported as ranges.
<sup>a</sup> Statistical tests performed: chi-square test of independence (for comparisons across sexes and sleep deprivation conditions); Wilcoxon rank-sum test (for age). <sup>b</sup> Statistics presented: Mean (SD). <sup>c</sup> Statistics presented: n (% of total)

The second dataset we examined is the Leipzig Study for Mind-Body-Emotion Interactions (LEMON), for which extensive details can be found elsewhere (Babayan et al., 2019). This cross-sectional study contains both young and older adults from Leipzig and the surrounding area. The final dataset we examined is the Stockholm SLEEPY Brain Study (Akerstedt, 2016). In this dataset, participants were instructed to maintain normal sleep patterns (sleep rested) or go to sleep 3 hours before waking up at their normal wake times (e.g., if a participant typically wakes up at 8am, for the sleep deprivation condition they would go to sleep at 5am but still wake up at 8am), to mimic real-world sleep deprivation.

Although the SLEEPY Brain study had two time points per participant (one sleep rested, one sleep deprived), only one time point included a diffusion-weighted scan, and therefore each participant was *either* in the sleep-rested or sleep-deprived condition, during diffusion scanning. Subject demographics in this study, with final N’s per dataset, are displayed in Table 1. We excluded subjects with poor quality diffusion-weighted scans, poor atlas registration, or missing data in statistical analyses. Poor scan and registration quality was determined through visual inspection. Tables 2-4 contain LC-MRI contrast sequence parameters, structural MRI parameters, and diffusion-weighted MRI parameters across studies, respectively.

**Table 2.**
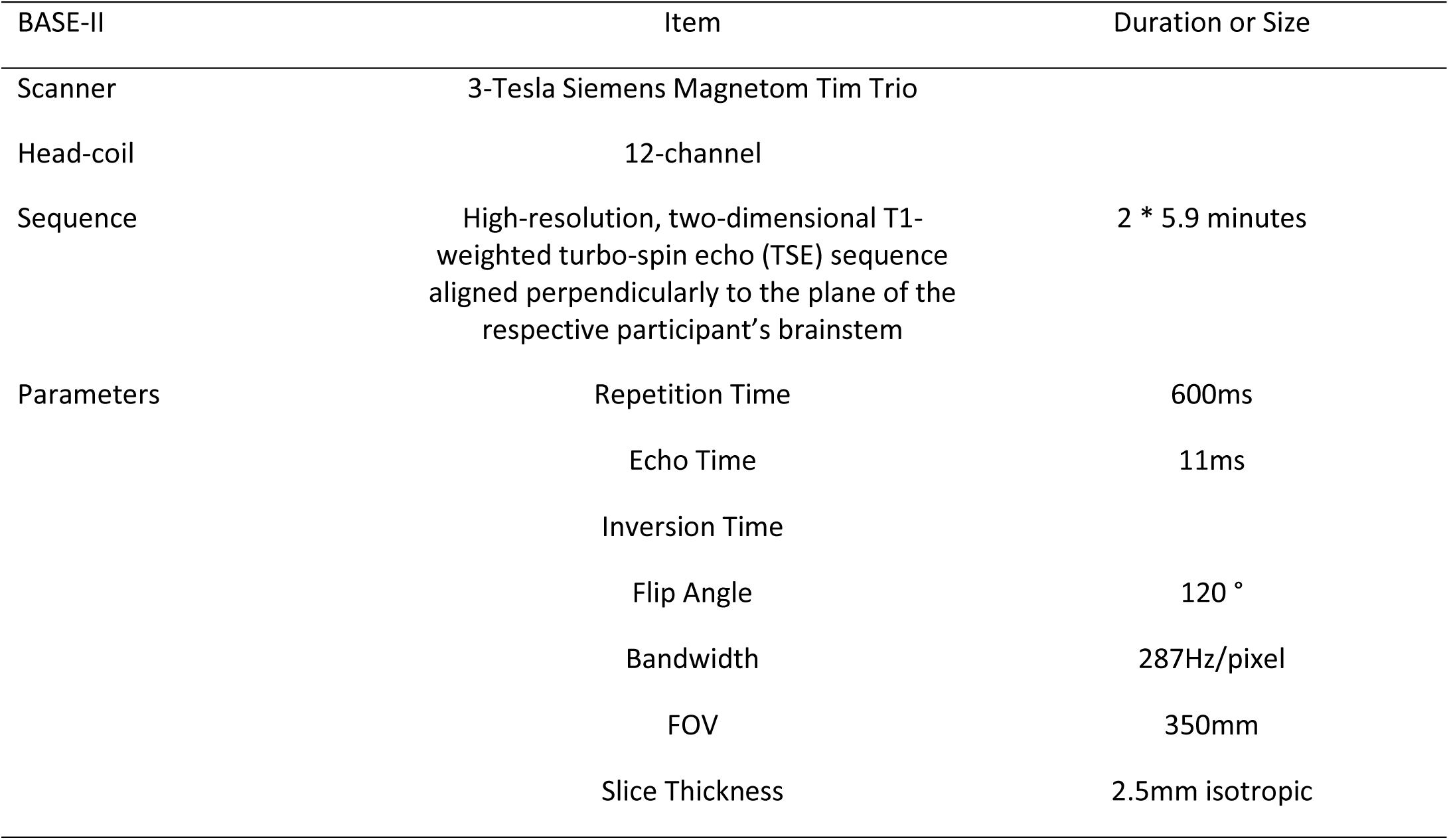
LC-MRI Contrast Sequence Parameters

### 2.1 DWI Processing

Using University of Southern California’s Laboratory of Neuroimaging (LONI) Pipeline, we applied FSL’s *(v6.3*) eddy-current correction, brain extraction tool, and resampling to isotropic resolution of 2mm^3^ (Dinov et al., 2009; Smith et al., 2004). We used MRtrix (*v3.1)* to compute fractional anisotropy (FA) and eigenvalue images (Tournier et al., 2019). Our atlas of the right and left LC nuclei was obtained from a LC meta-mask (Dahl et al., 2021) and the right and left noradrenergic bundles from Tang et al. (2020). As control regions, we utilized the previously defined right and left frontopontine tracts (Tang et al., 2018), which run along the ventral portion of the pons on either side of the basilar sulcus, terminating at the pontine nuclei. Figure 1 displays all three ROIs in MNI152 linear, 1mm resolution space.

**Figure 1.**
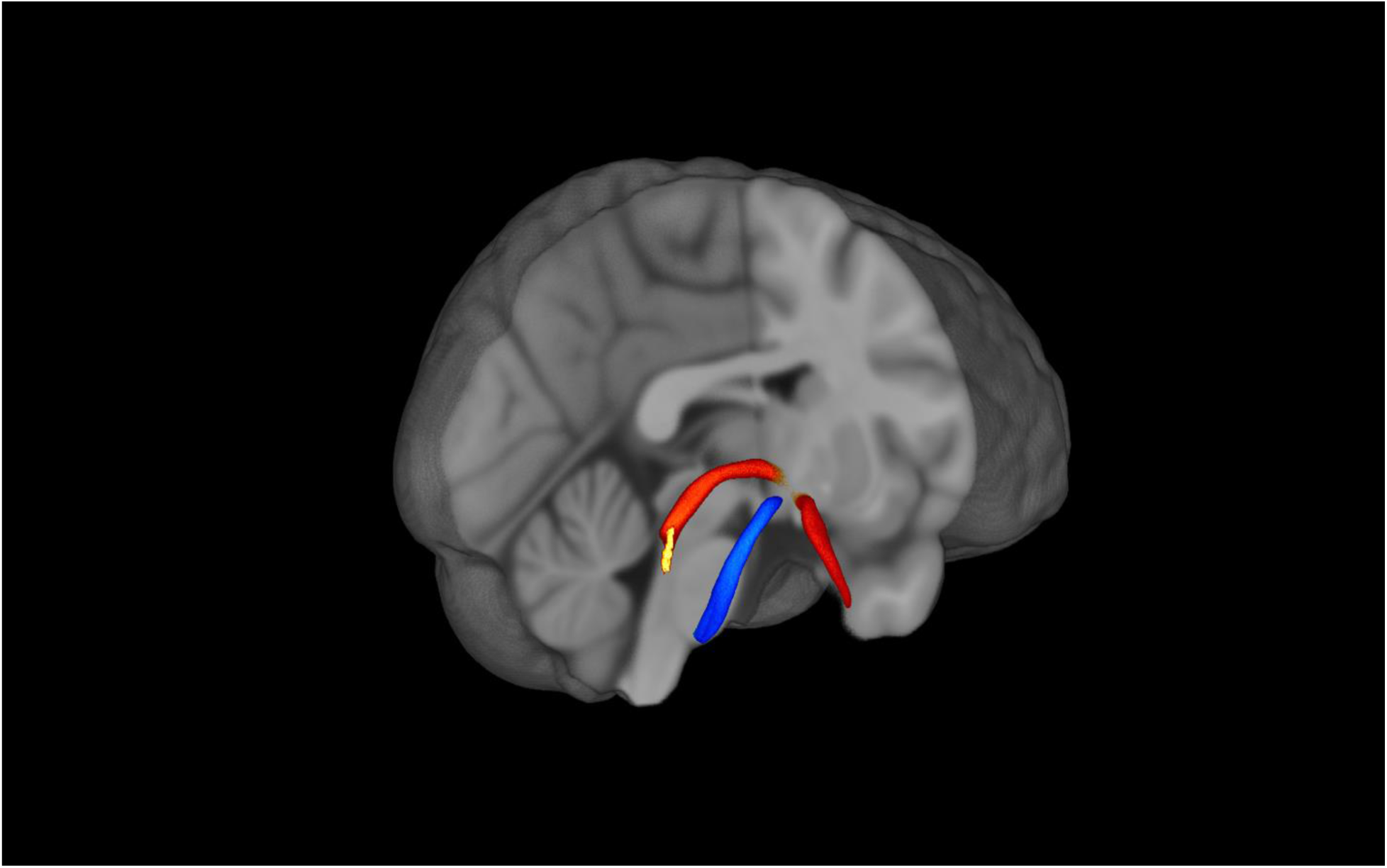
The ROI atlases of the Locus Coeruleus, Noradrenergic Bundle, and Frontopontine Tract. *Note.* Figure 1 displays the locus coeruleus (yellow), noradrenergic bundle (red), and frontopontine (blue) tracts registered to MNI152 space. The noradrenergic bundle is one continuous bundle (part of the temporal lobe segment is not pictured).

Both fractional anisotropy and atlas images were registered into MNI152 linear, 1mm brain space. Using ANTS nonlinear registration (Avants et al., 2008; Sun et al., 2020) the atlases were then backwarped into individual subject space with nearest neighbor interpolation. Registration quality was visualized using an in-house MATLAB script (*MATLAB ver. R2019a*). After accurate atlas registration to individual subject space was confirmed with visual inspection, mean and radial diffusion images were created from eigenvalue images in MATLAB with custom scripts. Atlases were then converted into a binary mask and multiplied by the diffusion image to provide fractional anisotropy, mean, and radial diffusivity values along the atlases, per voxel, within the native space. Diffusivity values were then averaged to provide one value per participant per ROI.

Since the noradrenergic bundle overlaps with a portion of the LC atlas, we conducted an along-tract analysis of fractional anisotropy of the noradrenergic bundle. 50 equidistant points were imposed along the noradrenergic bundle as discussed elsewhere (Sun et al., 2020). Each point was averaged across participants within younger or older adult groups. Anatomically, approximately the first 10 points represent most of the LC and points 30-50 represent areas of the entorhinal cortex. Fractional anisotropy at each point along the tract was compared between younger and older adults, shown in Figures 6-8.

### 2.2 Statistical Analyses

All statistical analyses were conducted using the R software (Team, 2020) with tidyverse and various additional packages (Ahlmann-Eltze, 2019; Aust & Barth, 2020; Kassambara; Lenth, 2021; Sjoberg et al., 2021; Wicham, 2017; Wickham, 2016; Xie, 2021). Correlation coefficients and 95% confidence intervals were used to identify the relationship between LC-MRI contrast and diffusivity measurements. Diffusivity and fractional anisotropy, mean diffusivity, and radial diffusivity values in the LC, ascending noradrenergic bundle, and frontopontine tract were analyzed within each dataset using a 2 × 2 × 3 × 2 factorial design in which age (younger, older) and gender (female, male) were between-subject factors and ROI (noradrenergic bundle, locus coeruleus, frontopontine tract) and hemisphere (left, right) were repeated-measures factors.

Levene’s tests were used to explore ANOVA assumptions of equal variances. We report effect sizes using 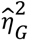 (generalized eta squared) for ANOVA effects and provide 95% confidence intervals to allow for comparisons across means. Greenhouse-Geisser correction was automatically computed for ANOVA departures from sphericity. For the along-tract analyses, *t*-tests were conducted for fractional anisotropy at each of the 50 equidistant points between younger and older adults. *P* values were false-discovery rate adjusted and surviving points of significant FA differences between age groups are plotted in Figures 6-8. Our focus was on fractional anisotropy but we include mean and radial diffusivity findings in the supplementary material. Lastly, to investigate if LC-FA diffusivity is associated with noradrenergic bundle-FA diffusivity, we conducted Pearson correlations and t-tests for each dataset.

## 3. Results

### 3.1 LC-MRI Contrasts and DTI

In the BASE-II dataset, there were no significant differences between young and older adults’ overall LC-MRI contrast values (Bachman et al., 2021; Dahl et al., 2019). We also did not observe significant associations between LC-MRI contrast and diffusivity in either the LC or ascending noradrenergic bundle. Correlation coefficients with FA and 95% confidence intervals for younger and older adults are displayed in Table 5 and Table 6, respectively. Previous studies have observed no overall age differences, but spatially confined age differences between caudal and rostral regions of the LC have been observed with LC-MRI contrast (Bachman et al., 2021; Dahl et al., 2019).

**Table 3.**
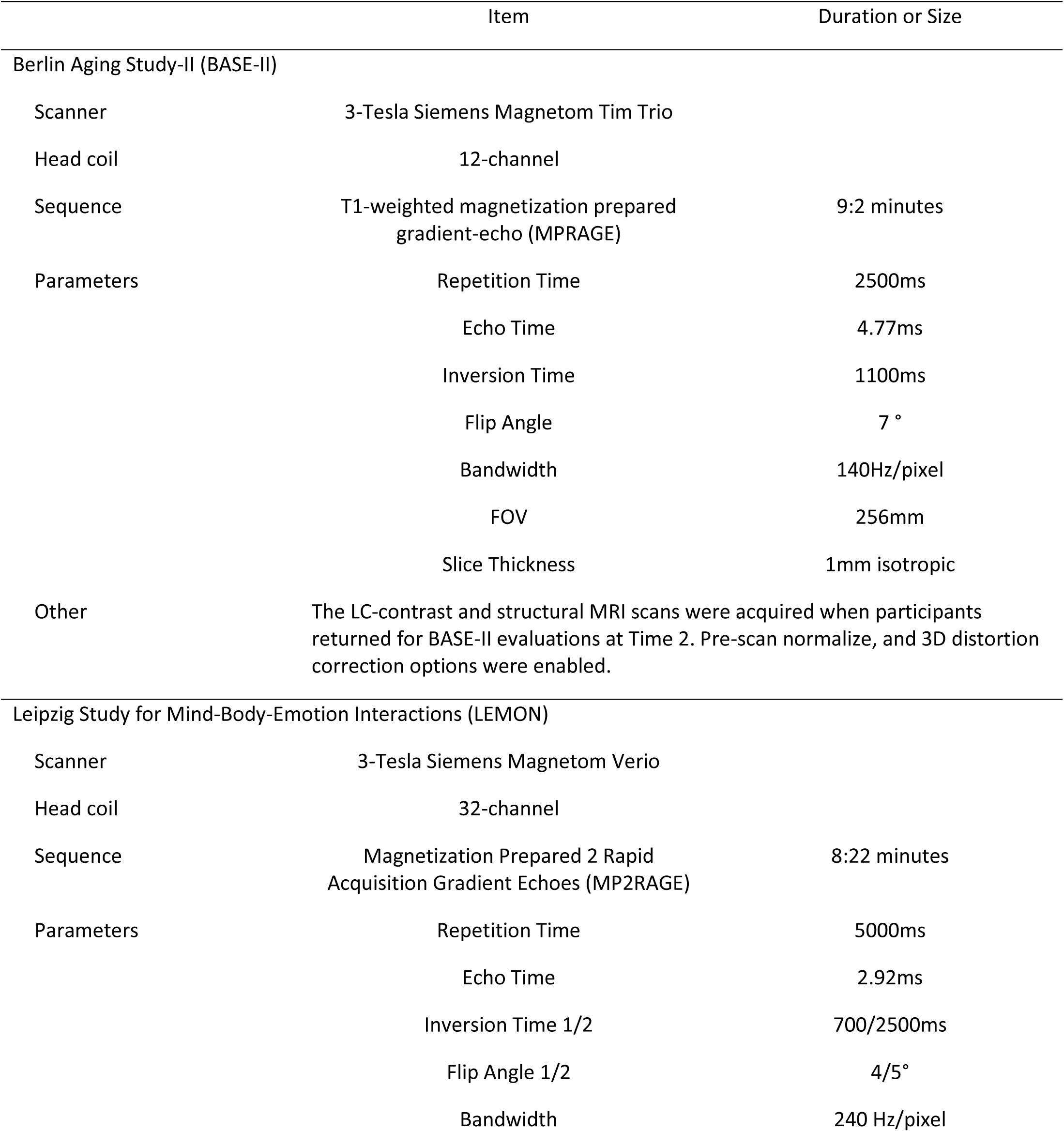
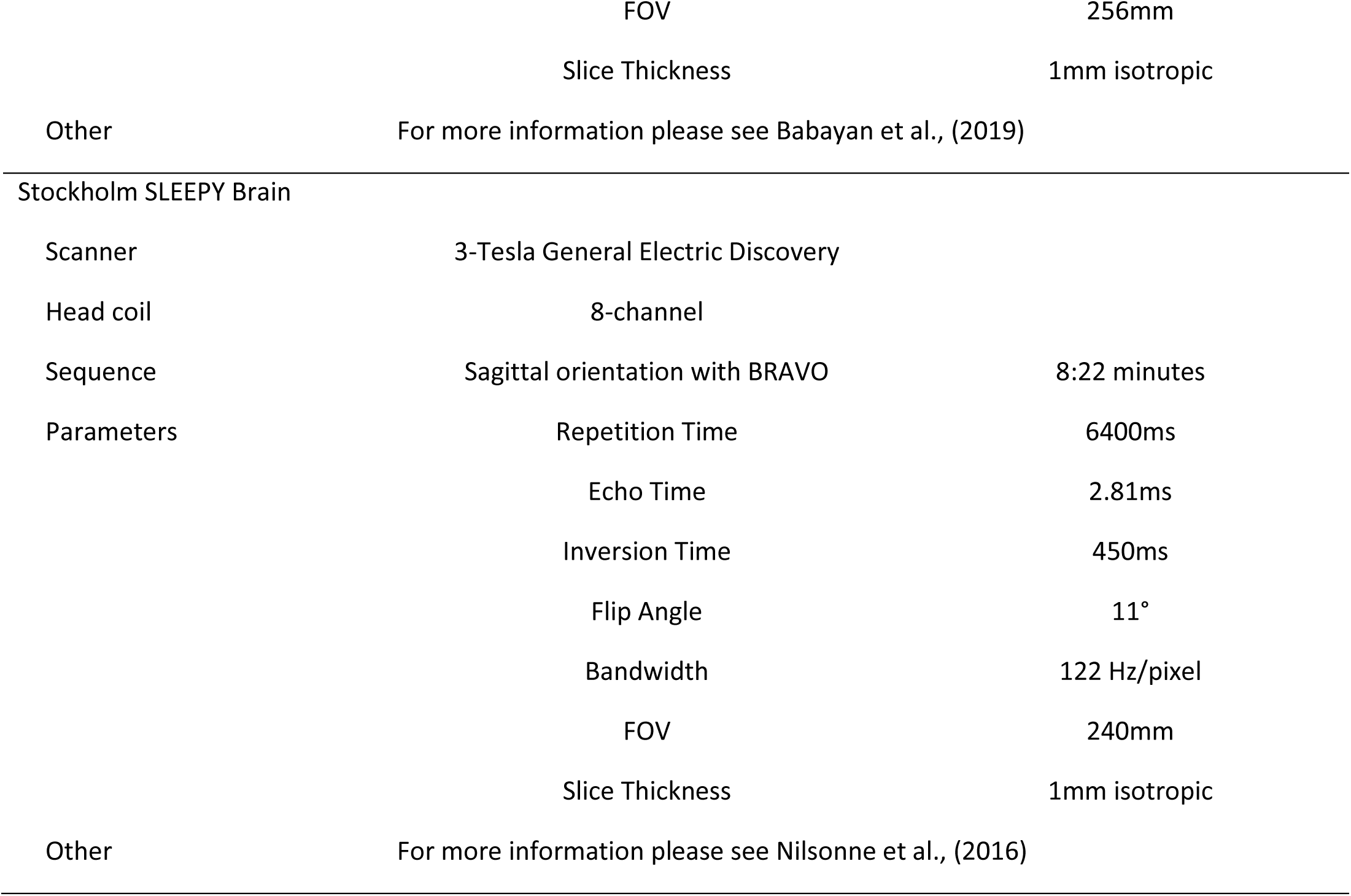
Structural MRI Sequence Parameters in Each Study

|  | Item | Duration or Size |
| --- | --- | --- |
| Berlin Aging Study-II (BASE-II) |  |  |
| Scanner | 3-Tesla Siemens Magnetom Tim Trio |  |
| Head coil | 12-channel |  |
| Sequence | T1-weighted magnetization prepared gradient-echo (MPRAGE) | 9:2 minutes |
| Parameters | Repetition Time | 2500ms |
|  | Echo Time | 4.77ms |
|  | Inversion Time | 1100ms |
|  | Flip Angle | 7 ° |
|  | Bandwidth | 140Hz/pixel |
|  | FOV | 256mm |
|  | Slice Thickness | 1mm isotropic |
| Other | The LC-contrast and structural MRI scans were acquired when participants returned for BASE-II evaluations at Time 2. Pre-scan normalize, and 3D distortion correction options were enabled. |  |
| Leipzig Study for Mind-Body-Emotion Interactions (LEMON) |  |  |
| Scanner | 3-Tesla Siemens Magnetom Verio |  |
| Head coil | 32-channel |  |
| Sequence | Magnetization Prepared 2 Rapid Acquisition Gradient Echoes (MP2RAGE) | 8:22 minutes |
| Parameters | Repetition Time | 5000ms |
|  | Echo Time | 2.92ms |
|  | Inversion Time 1/2 | 700/2500ms |
|  | Flip Angle 1/2 | 4/5° |
|  | Bandwidth | 240 Hz/pixel |
|  | FOV | 256mm |
|  | Slice Thickness | 1mm isotropic |
| Other | For more information please see Babayan et al., (2019) |  |
| Stockholm SLEEPY Brain |  |  |
| Scanner | 3-Tesla General Electric Discovery |  |
| Head coil | 8-channel |  |
| Sequence | Sagittal orientation with BRAVO | 8:22 minutes |
| Parameters | Repetition Time | 6400ms |
|  | Echo Time | 2.81ms |
|  | Inversion Time | 450ms |
|  | Flip Angle | 11° |
|  | Bandwidth | 122 Hz/pixel |
|  | FOV | 240mm |
|  | Slice Thickness | 1mm isotropic |
| Other | For more information please see Nilsonne et al., (2016) |  |

**Table 4.**
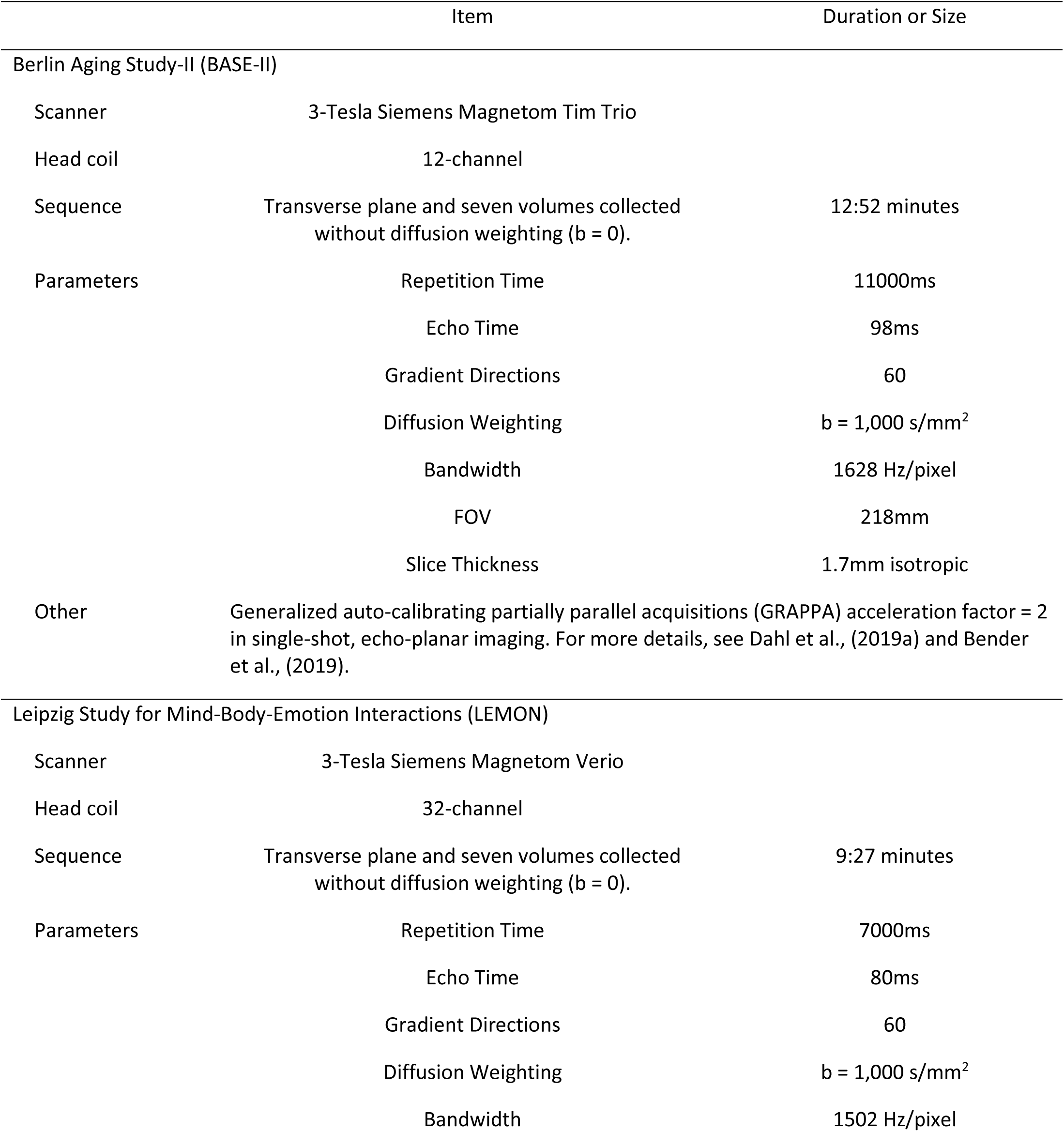
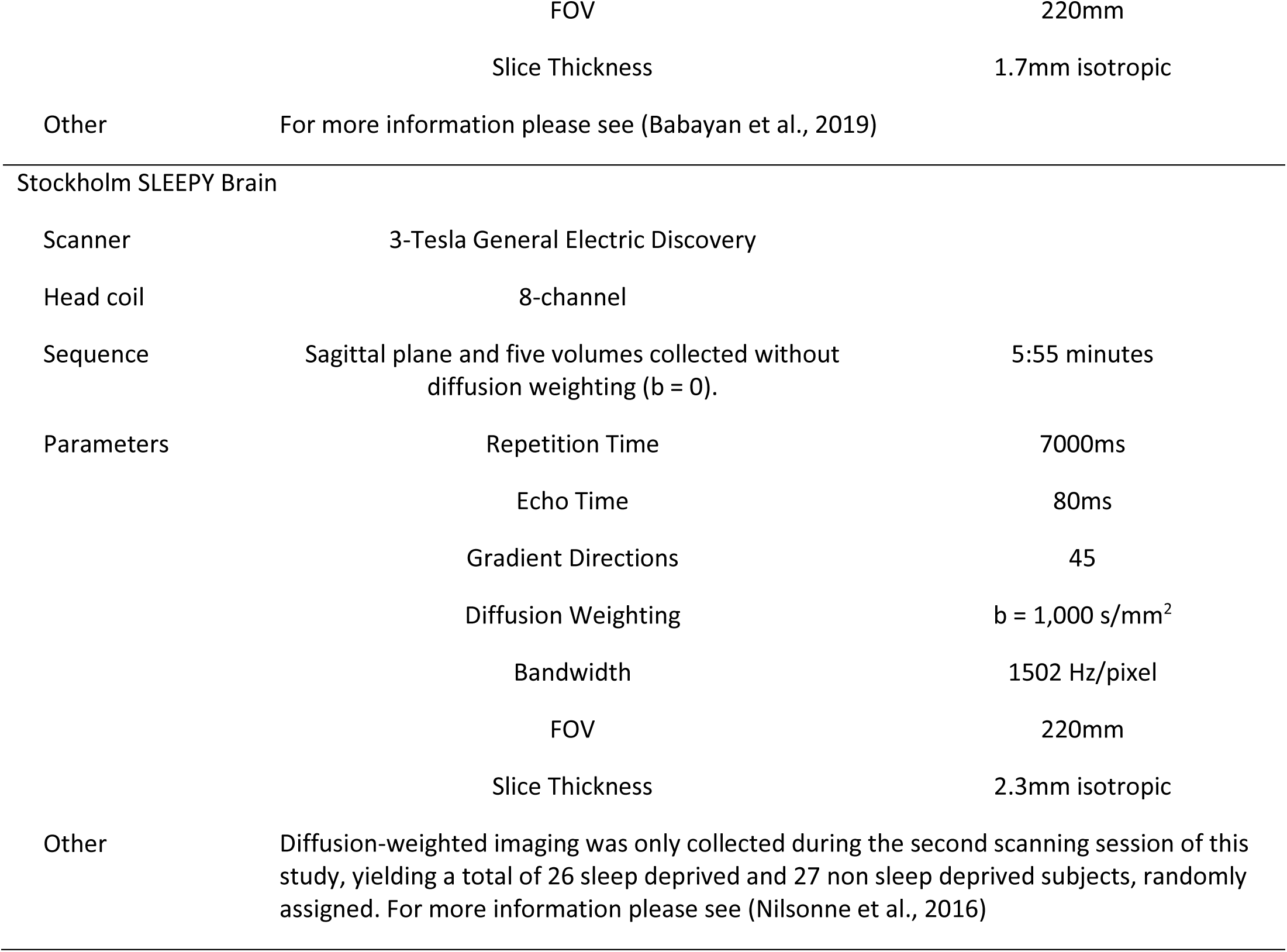
Diffusion MRI Sequence Parameters in Each Study

**Table 5.** Younger adults LC-MRI contrast correlations with confidence intervals

| Variable | LC-MRI Contrast |
| --- | --- |
| Noradrenergic bundle FA – Left hemisphere | -.07<br>[-.31, .18] |
| Noradrenergic bundle FA – Right hemisphere | -.09<br>[-.33, .16] |
| Locus Coeruleus FA – Left hemisphere | -.02<br>[-.27, .23] |
| Locus Coeruleus FA – Right hemisphere | -.06<br>[-.30, .19] |
| Frontopontine FA – Left hemisphere | .04<br>[-.21, .29] |
| Frontopontine FA – Right hemisphere | .15<br>[-.10, .39] |

*Note.* Values in square brackets indicate the 95% confidence interval for each correlation. The confidence interval is a plausible range of population correlations that could have caused the sample correlation (Cumming, 2014).

**Table 6.** Older adults LC-MRI contrast correlations with confidence intervals

| Variable | LC-MRI Contrast |
| --- | --- |
| Noradrenergic bundle FA – Left hemisphere | -.03<br>[-.15, .10] |
| Noradrenergic bundle FA – Right hemisphere | -.08<br>[-.20, .05] |
| Locus Coeruleus FA – Left hemisphere | .05<br>[-.08, .17] |
| Locus Coeruleus FA – Right hemisphere | -.10<br>[-.23, .02] |
| Frontopontine FA – Left hemisphere | -.06<br>[-.19, .06] |
| Frontopontine FA – Right hemisphere | -.02<br>[-.14, .11] |

*Note.* Values in square brackets indicate the 95% confidence interval for each correlation. The confidence interval is a plausible range of population correlations that could have caused the sample correlation (Cumming, 2014).

### 3.2 Fractional anisotropy in the LC is higher in older adults, relative to younger adults

Complete ANOVA tables for fractional anisotropy across datasets are displayed in Tables 7-9. Here in the text, we report the significant ANOVA interactions involving Age and ROI. In the BASE-II and LEMON datasets, we observed a significant interaction of Age x ROI for fractional anisotropy, *F*(1.57, 468.27) = 27.18, *p* < .001, 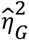 = .033, and *F*(1.79, 386.34) = 26.07, *p* < .001, 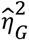 = 0.035, respectively (Table 7 and Table 8). We also observed a significant 3-way interaction of Age x ROI x Hemisphere for fractional anisotropy, *F*(1.62, 483.01) = 6.49, *p* = .003, 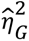 = .05, and *F*(1.63, 352.15) = 5.50, *p* = .008, 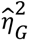 = 0.04, in the BASE-II and LEMON datasets, respectively.

**Table 7.** BASE-II Fractional Anisotropy Mixed ANOVA

| Variable | $F$ | $df_1^{GG}$ | $df_2^{GG}$ | $MSE$ | $p$ | $\hat{\eta}_G^2$ |
| --- | --- | --- | --- | --- | --- | --- |
| Age (Younger, Older) | 1.05 | 1 | 299 | 0.01 | .306 | .001 |
| Gender (Female, Male) | 7.33 | 1 | 299 | 0.01 | .007 | .007 |
| ROI (LC, Noradrenergic Bundle, Frontopontine) | 598.17 | 1.57 | 468.27 | 0.01 | <.001 | .426 |
| Hemisphere (Left, Right) | 125.52 | 1 | 299 | 0.00 | <.001 | .049 |
| Age × Gender | 1.21 | 1 | 299 | 0.01 | .273 | .001 |
| Age × ROI | 27.18 | 1.57 | 468.27 | 0.01 | <.001 | .033 |
| Gender × ROI | 1.97 | 1.57 | 468.27 | 0.01 | .151 | .002 |
| Age × Hemisphere | 0.00 | 1 | 299 | 0.00 | .980 | .000 |
| Gender × Hemisphere | 0.17 | 1 | 299 | 0.00 | .682 | .000 |
| ROI × Hemisphere | 73.81 | 1.62 | 483.01 | 0.00 | <.001 | .051 |
| Age × Gender × ROI | 0.46 | 1.57 | 468.27 | 0.01 | .584 | .001 |
| Age × Gender × Hemisphere | 0.46 | 1 | 299 | 0.00 | .497 | .000 |
| Age × ROI × Hemisphere | 6.49 | 1.62 | 483.01 | 0.00 | .003 | .005 |
| Gender × ROI × Hemisphere | 0.25 | 1.62 | 483.01 | 0.00 | .729 | .000 |
| Age × Gender × ROI × Hemisphere | 0.17 | 1.62 | 483.01 | 0.00 | .802 | .000 |

**Table 8.**
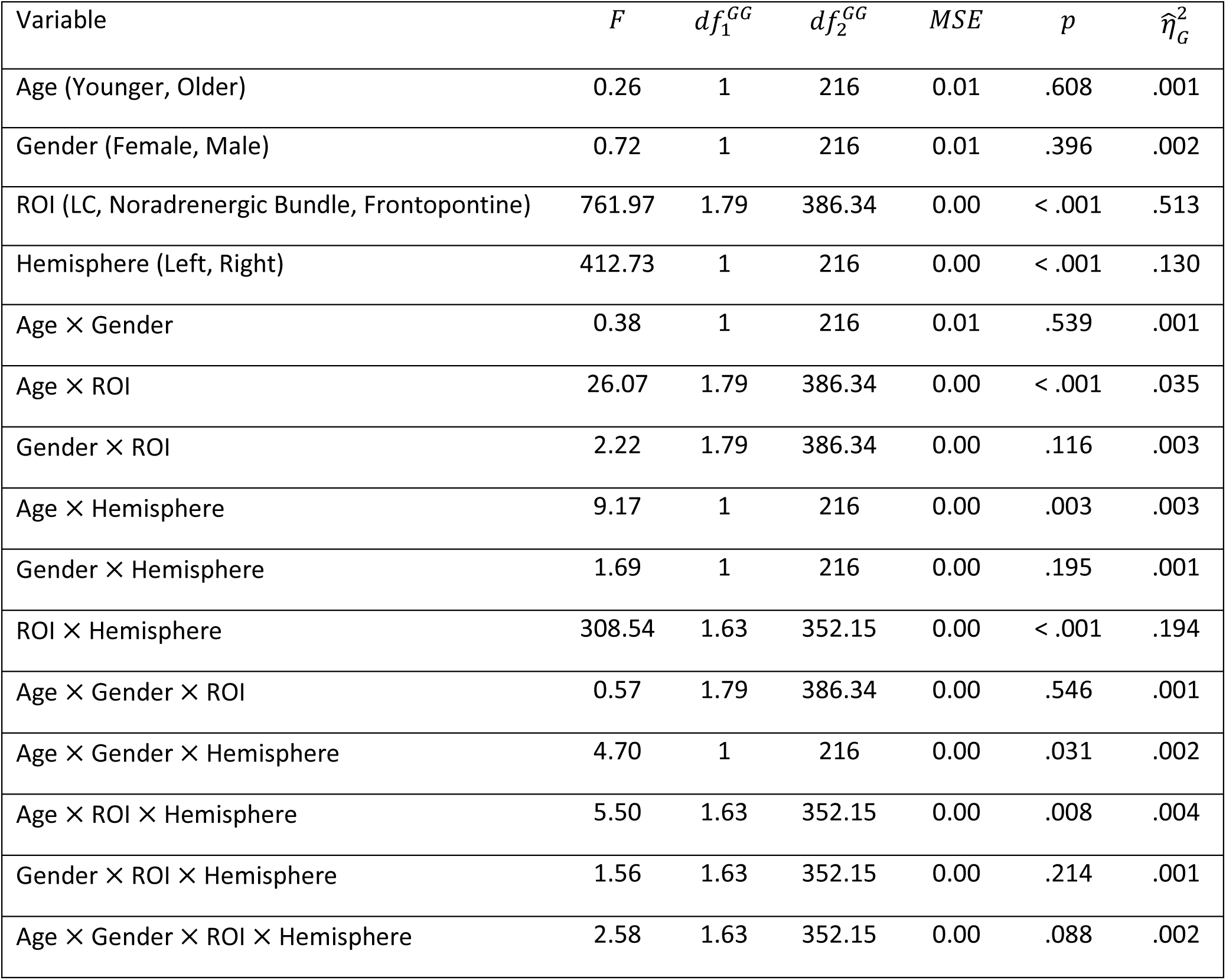
LEMON Fractional Anisotropy Mixed ANOVA

**Table 9.** SLEEPY Fractional Anisotropy Mixed ANOVA

| Variable | $F$ | $df_1^{GG}$ | $df_2^{GG}$ | $MSE$ | $p$ | $\hat{\eta}_G^2$ |
| --- | --- | --- | --- | --- | --- | --- |
| Age (Younger, Older) | 1.32 | 1 | 41 | 0.01 | .257 | .011 |
| Sleep Condition (Rested, Deprived) | 1.19 | 1 | 41 | 0.01 | .282 | .010 |
| ROI (LC, Noradrenergic Bundle, Frontopontine) | 115.99 | 1.76 | 72.17 | 0.00 | <.001 | .540 |
| Hemisphere (Left, Right) | 15.92 | 1 | 41 | 0.00 | <.001 | .040 |
| Age × Sleep Condition | 2.44 | 1 | 41 | 0.01 | .126 | .020 |
| Age × ROI | 3.09 | 1.76 | 72.17 | 0.00 | .058 | .030 |
| Sleep Condition × ROI | 2.48 | 1.76 | 72.17 | 0.00 | .098 | .024 |
| Age × Hemisphere | 1.27 | 1 | 41 | 0.00 | .267 | .003 |
| Sleep Condition × Hemisphere | 1.22 | 1 | 41 | 0.00 | .275 | .003 |
| ROI × Hemisphere | 18.37 | 1.57 | 64.45 | 0.00 | <.001 | .054 |
| Age × Sleep Condition × ROI | 3.12 | 1.76 | 72.17 | 0.00 | .056 | .031 |
| Age × Sleep Condition × Hemisphere | 2.08 | 1 | 41 | 0.00 | .157 | .005 |
| Age × ROI × Hemisphere | 0.34 | 1.57 | 64.45 | 0.00 | .659 | .001 |
| Sleep Condition × ROI × Hemisphere | 0.24 | 1.57 | 64.45 | 0.00 | .732 | .001 |
| Age × Sleep Condition × ROI × Hemisphere | 0.24 | 1.57 | 64.45 | 0.00 | .731 | .001 |

Tables 10 and 11 report means and 95% confidence intervals for fractional anisotropy for each ROI between age groups, in each hemisphere. We observed significantly less fractional anisotropy in the LC and significantly more fractional anisotropy in the noradrenergic bundle of younger adults compared to older adults, in both the BASE-II and LEMON datasets (Tables 10 and 11; Figures 2 and 3). We observed no significant differences in frontopontine tract fractional anisotropy between younger and older adults in either BASE-II or LEMON datasets.

**Figure 2.**
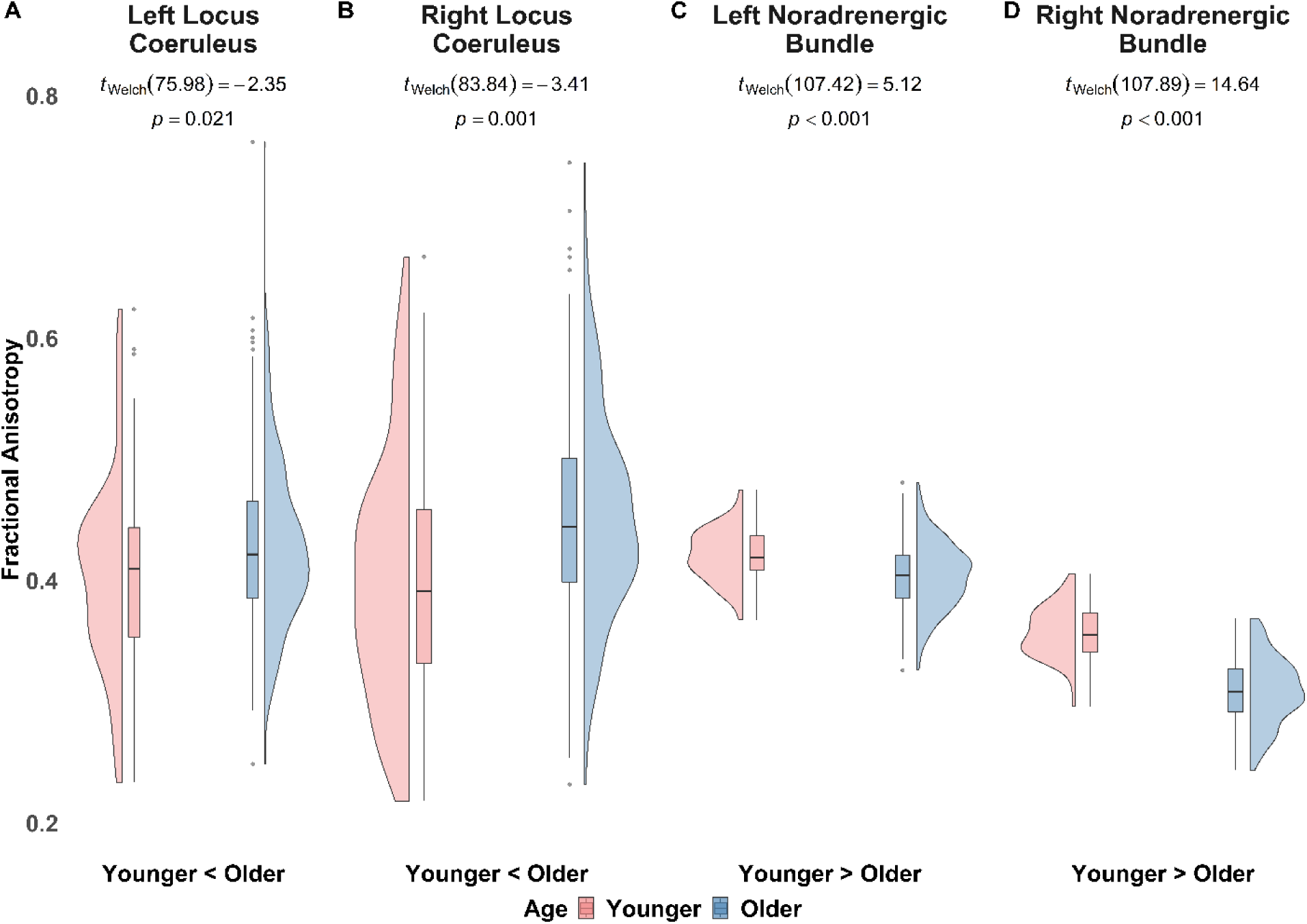
BASE-II Fractional Anisotropy in Left and Right Locus Coeruleus and Noradrenergic Bundles in Younger and Older Adults. *Note.* Figure 2 displays fractional anisotropy between younger and older adults from the BASE-II cohort. In the left locus coeruleus (A) and right locus coeruleus (B), we observed lower fractional anisotropy in younger adults, compared to older adults. In the left noradrenergic bundle (C) and right noradrenergic bundle (D) we observed higher fractional anisotropy in younger adults, relative to older adults.

**Figure 3.**
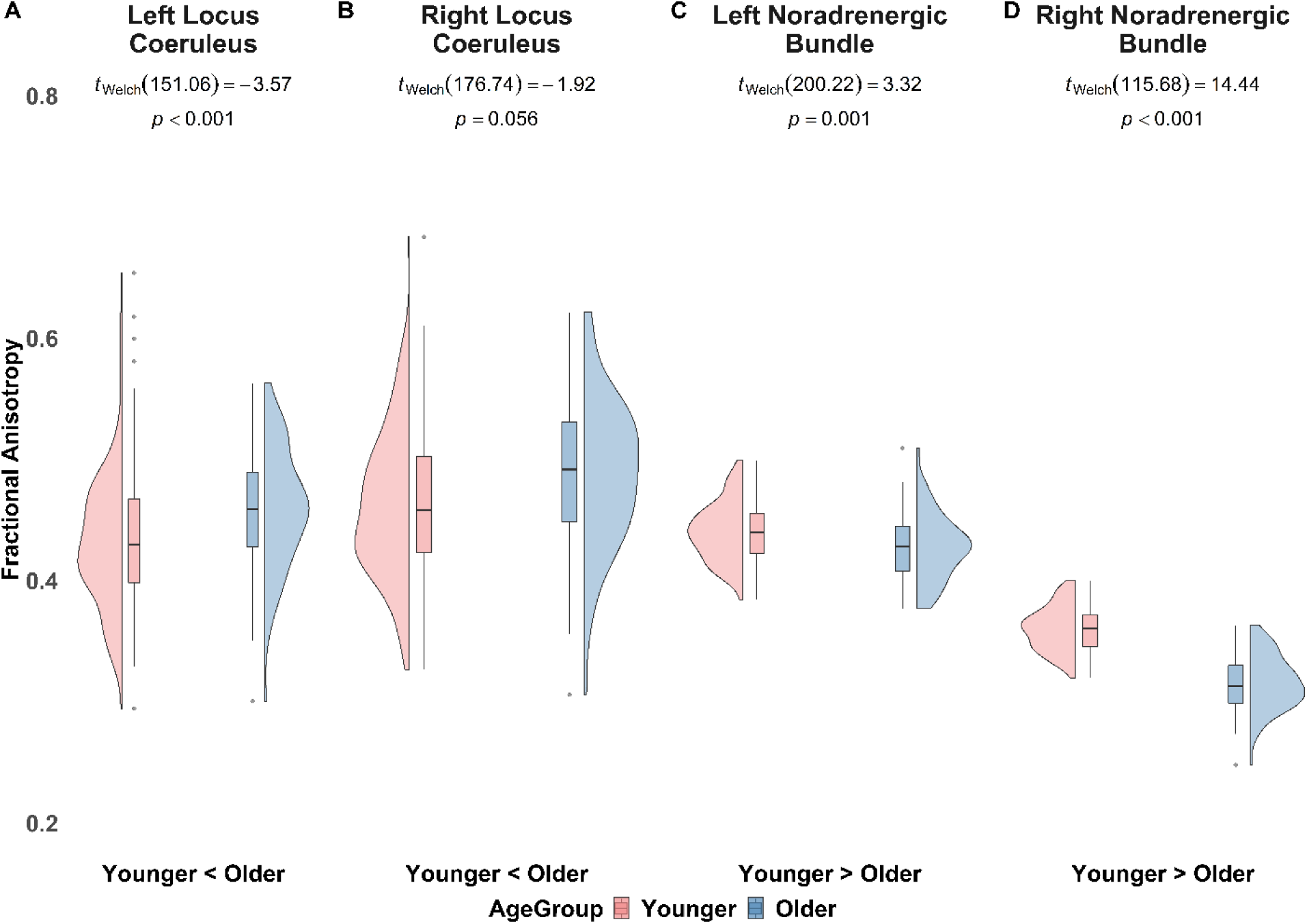
LEMON Fractional Anisotropy in Left and Right Locus Coeruleus and Noradrenergic Bundles in Younger and Older Adults. *Note.* Figure 3 displays our BASE-II replicated fractional anisotropy findings between younger and older adults in the LEMON dataset. In the left locus coeruleus (A) and right locus coeruleus (B), we observed lower fractional anisotropy in younger adults, compared to older adults. In the left noradrenergic bundle (C) and right noradrenergic bundle (D) we observed higher fractional anisotropy in younger adults, relative to older adults.

**Table 10.** BASE-II Fractional Anisotropy Means, Standard Error, Degrees of Freedom and 95% Confidence Intervals

| BASE-II | Locus Coeruleus |  | Noradrenergic Bundle |  | Frontopontine Tract |  |
| --- | --- | --- | --- | --- | --- | --- |
|  | Young Adult | Older Adult | Young Adult | Older Adult | Young Adult | Older Adult |
| Left Hemisphere |  |  |  |  |  |  |
| $M^a$ | 0.396 | 0.426 | 0.421 | 0.403 | 0.563 | 0.567 |
| $SE$ | 0.01 | 0.005 | 0.004 | 0.002 | 0.008 | 0.004 |
| Lower CI | 0.377 | 0.417 | 0.413 | 0.4 | 0.547 | 0.559 |
| Upper CI | 0.416 | 0.436 | 0.428 | 0.407 | 0.58 | 0.574 |
| Right Hemisphere |  |  |  |  |  |  |
| $M^a$ | 0.397 | 0.452 | 0.356 | 0.309 | 0.515 | 0.522 |
| $SE$ | 0.014 | 0.007 | 0.004 | 0.002 | 0.008 | 0.004 |
| Lower CI | 0.369 | 0.439 | 0.348 | 0.306 | 0.5 | 0.515 |
| Upper CI | 0.424 | 0.465 | 0.363 | 0.313 | 0.53 | 0.529 |

*Note.* CI = confidence interval<sup>a</sup> degrees of freedom = 299.

**Table 11.** LEMON Fractional Anisotropy Means, Standard Error, Degrees of Freedom and 95% Confidence Intervals

| LEMON | Locus Coeruleus |  | Noradrenergic Bundle |  | Frontopontine Tract |  |
| --- | --- | --- | --- | --- | --- | --- |
|  | Young Adult | Older Adult | Young Adult | Older Adult | Young Adult | Older Adult |
| Left Hemisphere |  |  |  |  |  |  |
| *M*<sup>a</sup> | 0.427 | 0.459 | 0.445 | 0.427 | 0.584 | 0.582 |
| *SE* | 0.005 | 0.006 | 0.003 | 0.005 | 0.006 | 0.008 |
| Lower CI | 0.418 | 0.446 | 0.438 | 0.418 | 0.572 | 0.567 |
| Upper CI | 0.437 | 0.472 | 0.451 | 0.436 | 0.595 | 0.597 |
| Right Hemisphere |  |  |  |  |  |  |
| *M*<sup>a</sup> | 0.473 | 0.487 | 0.365 | 0.315 | 0.497 | 0.503 |
| *SE* | 0.007 | 0.009 | 0.003 | 0.004 | 0.005 | 0.007 |
| Lower CI | 0.46 | 0.47 | 0.359 | 0.308 | 0.487 | 0.489 |
| Upper CI | 0.486 | 0.505 | 0.37 | 0.323 | 0.508 | 0.517 |

*Note.* CI = confidence interval<sup>a</sup> degrees of freedom = 216.

For the SLEEPY Brain dataset, we used the same ANOVA factorial design as in the other datasets with sleep condition (rested, deprived) as an additional between-subjects factor. The Age x ROI interaction did not quite reach significance, *F*(1.76, 72.17) = 3.09, *p* = .058, 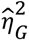 = 0.03, (Table 9). Likewise, the Age x ROI x Sleep Condition interaction did not quite achieve significance, *F*(1.76, 72.17) = 3.12, *p* = .056, 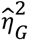 = 0.031. Here, we report planned contrasts of fractional anisotropy across ROIs between age groups and sleep conditions. Means and confidence intervals for the SLEEPY dataset are available in Table 12, and plots for between and within age group differences are shown in Figures 4 and 5, respectively.

**Figure 4.**
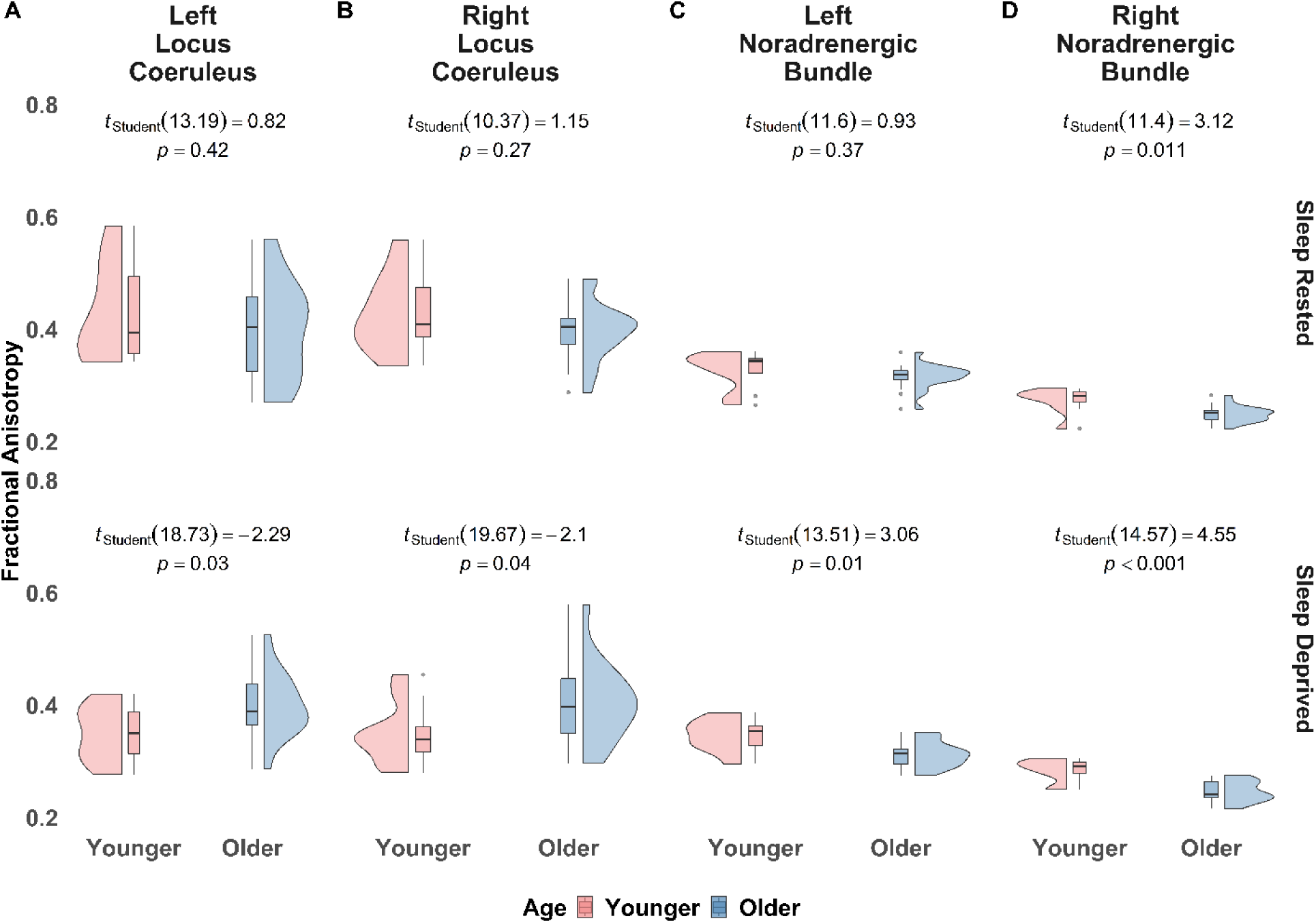
SLEEPY Brain Fractional Anisotropy in Left and Right Locus Coeruleus and Noradrenergic Bundles in Younger and Older Adults. *Note.* Figure 4 displays fractional anisotropy between younger and older adults in either the sleep rested or sleep deprived conditions. In the left (A) and right (B) locus coeruleus, we observed no significant differences in fractional anisotropy between young and older adults in the sleep rested condition. However, in the sleep deprived condition, younger adults displayed significantly lower fractional anisotropy, compared to older adults. In the left noradrenergic bundle (C) we observed no significant differences between young and older adult fractional anisotropy in the sleep rested condition. In the sleep deprived condition (C – row 2), younger adults had significantly higher fractional anisotropy, relative to older adults. In the right noradrenergic bundle (D) younger adults, compared to older adults, had significantly higher fractional anisotropy in the sleep rested condition. Fractional anisotropy differences were more pronounced in the sleep deprived condition.

**Figure 5.**
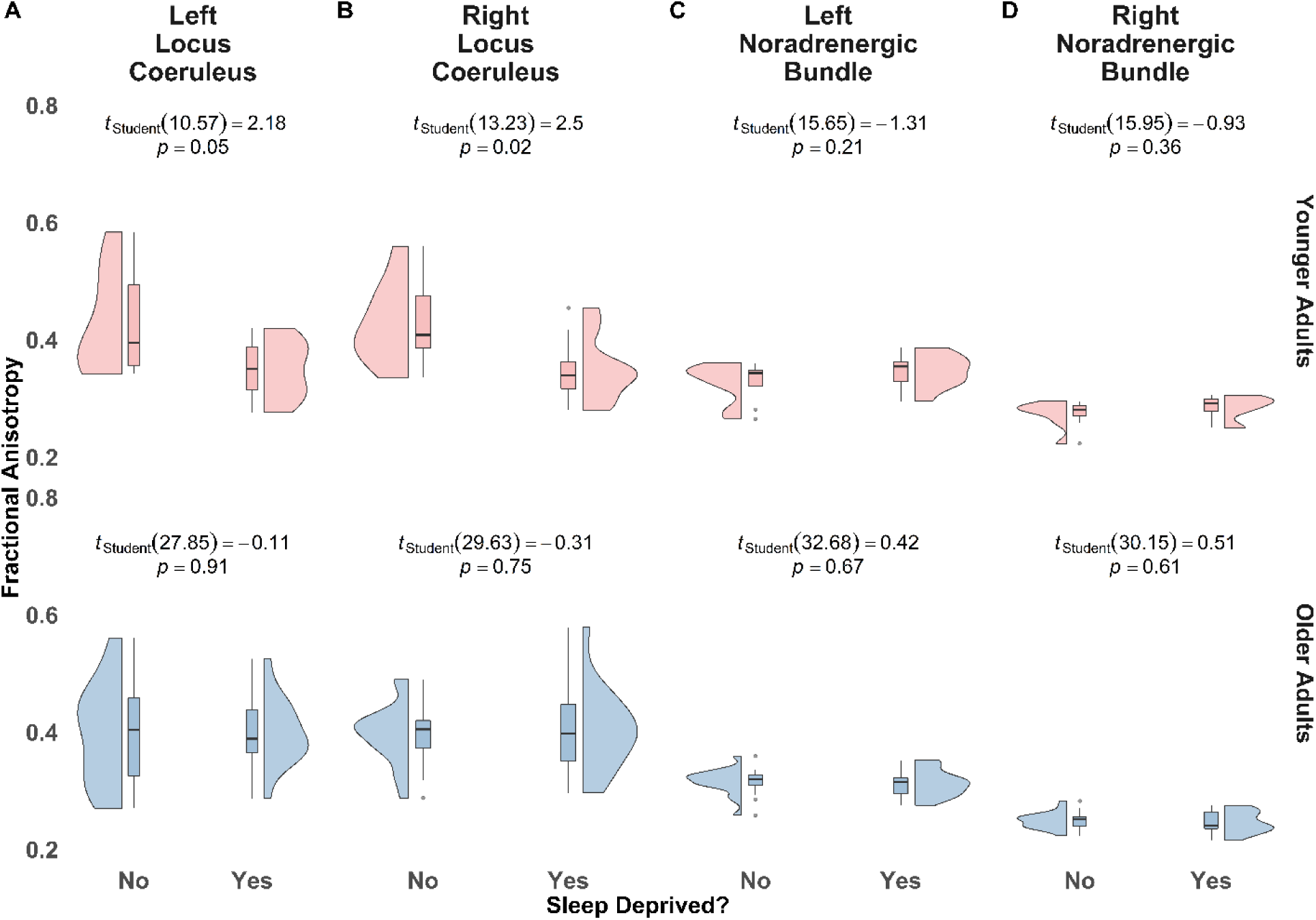
SLEEPY Brain Fractional Anisotropy in Left and Right Locus Coeruleus and Noradrenergic Bundles between Younger and Older Adults. *Note.* Figure 5 displays fractional anisotropy between sleep rested or sleep deprived young and older adults. In the left locus coeruleus (A) and right locus coeruleus (B) we observed significantly higher fractional anisotropy in sleep rested young adults, compared to sleep deprived young adults. We observed no significant difference of fractional anisotropy between sleep rested and sleep deprived older adults in the left or right locus coeruleus. In the left noradrenergic bundle (C) and right noradrenergic bundle (D) we observed no significant difference in fractional anisotropy in sleep rested or sleep deprived younger adults; nor sleep rested or sleep deprived older adults.

**Table 12.** SLEEPY Fractional Anisotropy Means, Standard Error, Degrees of Freedom and 95% Confidence Intervals

| SLEEPY | Locus Coeruleus |  | Noradrenergic Bundle |  | Frontopontine Tract |  |
| --- | --- | --- | --- | --- | --- | --- |
|  | Young Adult | Older Adult | Young Adult | Older Adult | Young Adult | Older Adult |
| Left Hemisphere, Rested |  |  |  |  |  |  |
| *M*<sup>a</sup> | 0.405 | 0.399 | 0.323 | 0.315 | 0.475 | 0.446 |
| *SE* | 0.027 | 0.019 | 0.01 | 0.007 | 0.029 | 0.02 |
| Lower CI | 0.35 | 0.36 | 0.303 | 0.301 | 0.417 | 0.405 |
| Upper CI | 0.46 | 0.437 | 0.344 | 0.33 | 0.533 | 0.486 |
| Left Hemisphere, Deprived |  |  |  |  |  |  |
| *M*<sup>a</sup> | 0.343 | 0.395 | 0.35 | 0.311 | 0.469 | 0.463 |
| *SE* | 0.025 | 0.018 | 0.009 | 0.007 | 0.027 | 0.019 |
| Lower CI | 0.292 | 0.358 | 0.331 | 0.298 | 0.415 | 0.424 |
| Upper CI | 0.394 | 0.432 | 0.369 | 0.325 | 0.523 | 0.503 |
| Right Hemisphere, Rested |  |  |  |  |  |  |
| *M*<sup>a</sup> | 0.435 | 0.394 | 0.272 | 0.249 | 0.49 | 0.415 |
| *SE* | 0.023 | 0.016 | 0.007 | 0.005 | 0.028 | 0.02 |
| Lower CI | 0.389 | 0.361 | 0.257 | 0.239 | 0.433 | 0.375 |
| Upper CI | 0.482 | 0.427 | 0.287 | 0.26 | 0.547 | 0.455 |
| Right Hemisphere, Deprived |  |  |  |  |  |  |
| *M*<sup>a</sup> | 0.339 | 0.404 | 0.282 | 0.246 | 0.44 | 0.43 |
| *SE* | 0.021 | 0.016 | 0.007 | 0.005 | 0.026 | 0.019 |
| Lower CI | 0.295 | 0.372 | 0.268 | 0.236 | 0.387 | 0.392 |
| Upper CI | 0.382 | 0.436 | 0.296 | 0.256 | 0.493 | 0.469 |

*Note.* CI = confidence interval.<sup>a</sup> degrees of freedom = 41.

In the *sleep rested* condition, we observed no significant differences in fractional anisotropy between young and older adults in the left or right LC, *t*(41) = 0.181, *p* = .857, and *t*(41) = 1.469*, p* = .149, respectively. We observed no significant differences in fractional anisotropy between *sleep rested* young and older adults in the left noradrenergic bundle, *t*(41) = .619, *p* = .539, but significantly higher fractional anisotropy in the right noradrenergic bundle of *sleep rested* younger adults, compared to older, *t*(41) = 2.56, *p* = .014. Similarly, we observed no significant differences in fractional anisotropy between *sleep rested* young and older adults in the left frontopontine tract, *t*(41) = 0.838, *p* = .407, but significantly higher fractional anisotropy in the right frontopontine tract of *sleep rested* younger adults, compared to older, *t*(41) = 2.161, *p* = .036.

Within the *sleep deprived* condition, younger adults had lower fractional anisotropy than older adults in the LC that did not quite achieve significance in the left locus coeruleus, *t*(41) = -1.658, *p* = .104, but was significant in the right locus coeruleus, *t*(41) = -2.451, *p* = .018. We also observed significantly higher fractional anisotropy in *sleep deprived* younger adults, compared with older adults, in both the left and right noradrenergic bundle, *t*(41) = 3.286, *p* = .002 and *t*(41) = 4.291, *p* < .001, respectively. No significant differences were observed between younger and older adult’s fractional anisotropy in the *sleep deprived* conditions in the left or right frontopontine tract, *t*(41) = 0.163, *p* = .870, and *t*(41) = 0.296, *p* = .769, respectively.

The BASE-II and LEMON along-tract analyses (Figure 6-7) show effects that are consistent with the LC and noradrenergic bundle results described above. Along the first 10 points, which approximately represent regions close to the LC, younger adults display significantly lower fractional anisotropy, relative to older adults. In the remaining tract points, younger adults had higher fractional anisotropy, relative to older adults, with significant age differences toward the end of the tract, in the region of the entorhinal cortex. We did not observe significant differences along the noradrenergic bundle between younger and older adults in the SLEEPY dataset, to the degree observed in the BASE-II and LEMON datasets.

**Figure 6.**
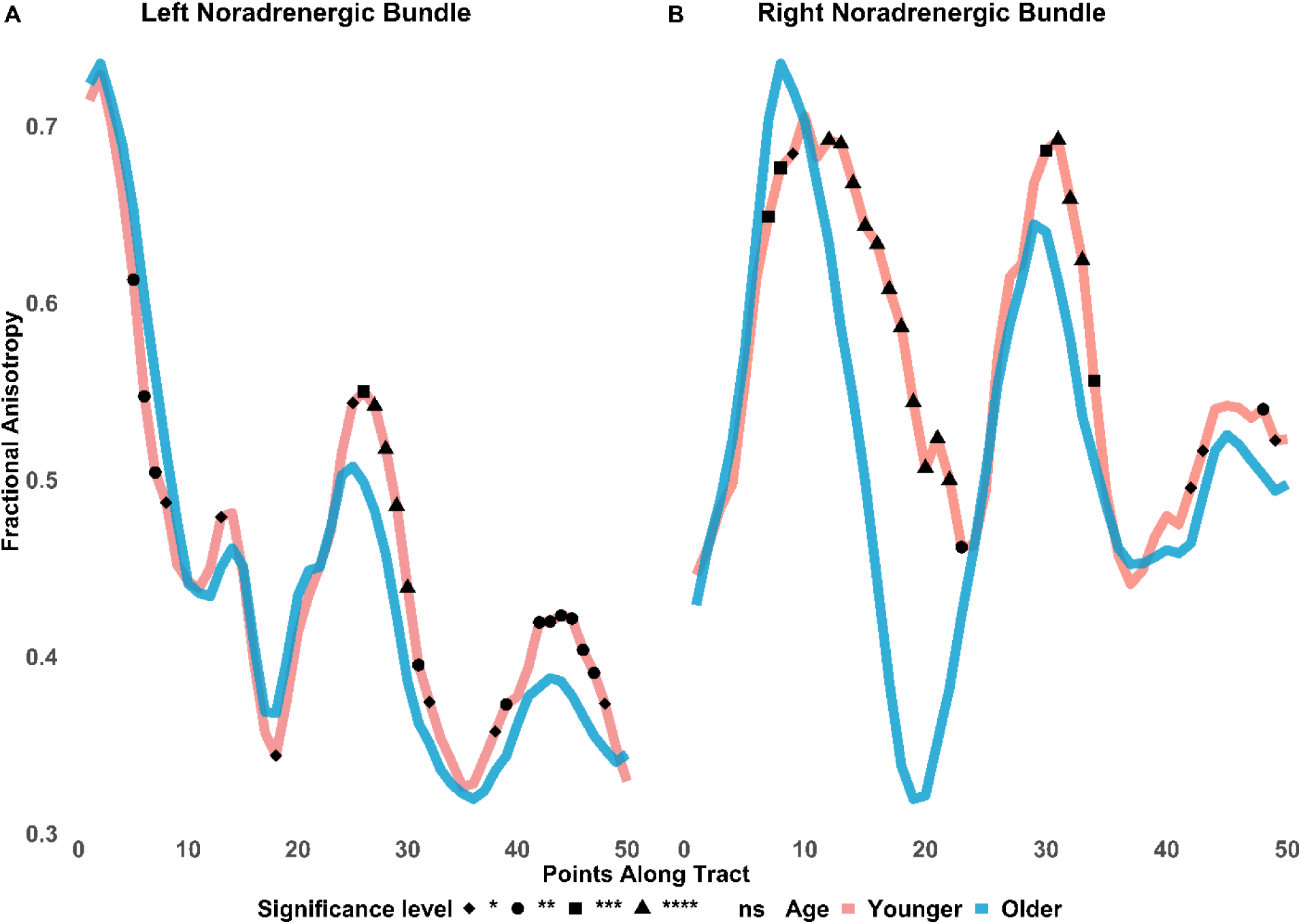
BASE-II Fractional Anisotropy Along the Noradrenergic Bundle. *Note.* ns = not significant; ns not assigned shape. Fractional anisotropy differences between younger and older adults are shown along the noradrenergic bundle. The bundle was divided into 50 equidistant points and mean fractional anisotropy was calculated for each age group at each point. Younger adults had significantly lower fractional anisotropy in the first 10 points of the noradrenergic bundles which would correspond to the area of the locus coeruleus. In contrast, around the entorhinal cortex, younger adults show higher fractional anisotropy, compared to older adults. * p ≤ .05. ** p ≤ .01. *** p ≤ .001. **** p ≤ .0001. FDR adjusted.

**Figure 7.**
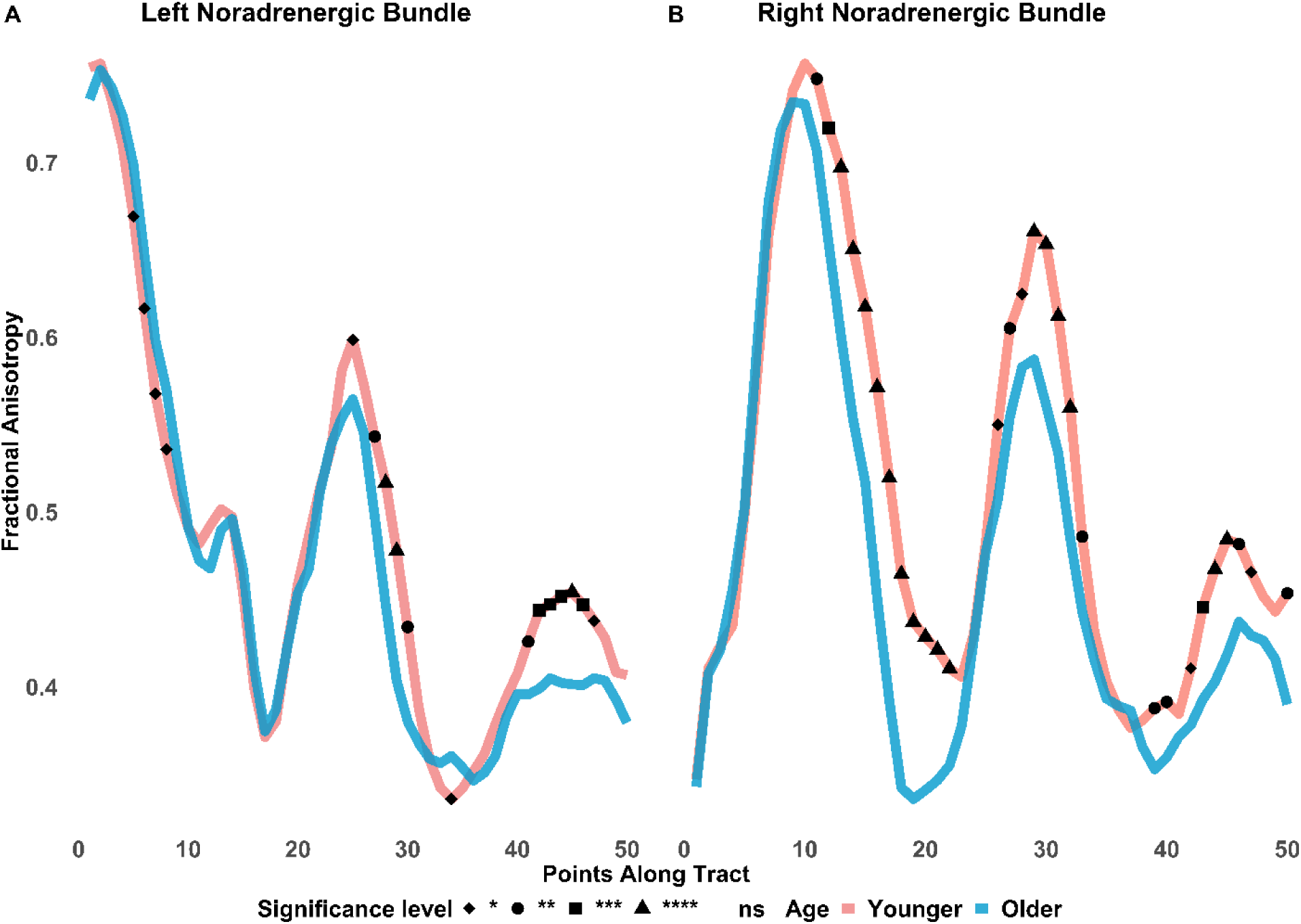
LEMON Fractional Anisotropy Along the Noradrenergic Bundle. *Note.* ns = not significant; ns not assigned shape. Fractional anisotropy differences between LEMON younger and older adults are shown along the noradrenergic bundle. The bundle was divided into 50 equidistant points and mean fractional anisotropy was calculated for each age group at each point. Younger adults had significantly lower fractional anisotropy in the first 10 points of the noradrenergic bundle, more so in the left than right, which would correspond to the area of the locus coeruleus. In contrast, younger adults showed higher fractional anisotropy and significantly greater differences around the entorhinal cortex, compared to older adults. * p ≤ .05. ** p ≤ .01. *** p ≤ .001. **** p ≤ .0001. FDR adjusted.

**Figure 8.**
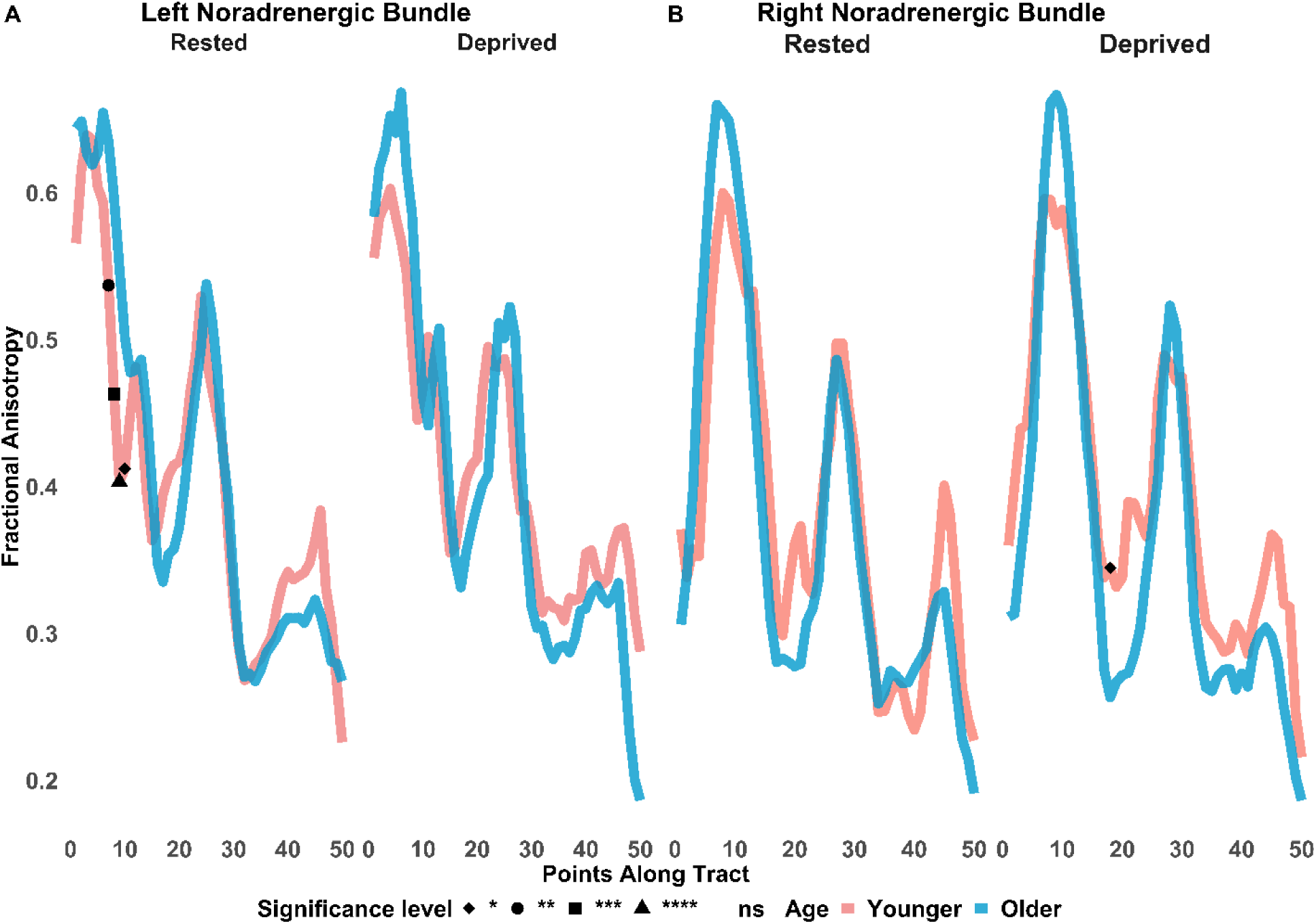
SLEEPY Fractional Anisotropy Along the Noradrenergic Bundle between Sleep Rested and Sleep Deprived Younger or Older Adults. *Note.* ns = not significant; ns not assigned shape. Fractional anisotropy differences between SLEEPY younger and older adults, either sleep rested or sleep deprived, are shown along the noradrenergic bundle. The bundle was divided into 50 equidistant points and mean fractional anisotropy was calculated for each age group at each point. Younger adults display significantly lower fractional anisotropy in the first 10 points (reflecting the locus coeruleus) of the left noradrenergic bundle, when sleep rested. * p ≤ .05. ** p ≤ .01. *** p ≤ .001. **** p ≤ .0001. FDR adjusted.

Finally, we observed significant positive correlations between LC fractional anisotropy and noradrenergic bundle fractional anisotropy within the BASE-II older adult cohort in the left and right hemispheres, *r*(243) = .24, *p* < .001 and *r*(243) = .19, *p* < .001, respectively. However, only three percent of the variance was accounted for (R^2^_adj_ = 0.03). We did not observe any significant relationship in the BASE-II young adult cohort. We were also not able to replicate these findings in the LEMON or SLEEPY datasets.

## 4. Discussion

Fractional anisotropy has been well established to correlate with white matter integrity, increasing until the age of about 35-40 and decreasing into late life or with disease (Kochunov et al., 2012; Li et al., 2016). Additionally, mean, and radial diffusivity are typically negatively correlated with fractional anisotropy (Beaudet et al., 2020; Li et al., 2016). Here, using three publicly available datasets (Akerstedt, 2016; Babayan et al., 2019; Delius et al., 2015), we examined the age-related diffusivity of the LC, ascending noradrenergic bundle, and, as a control, frontopontine white matter tracts. We replicated Langley et al.’s (2020) findings of higher fractional anisotropy in the LC in older adults compared with younger adults, across two large datasets (BASE-II; LEMON).

Interestingly, we did not observe these differences in the locus coeruleus in the SLEEPY Brain dataset, between young and older adults, who were *sleep rested*, but only those who were *sleep deprived*. We did, however, observe typical age differences in diffusivity of the noradrenergic bundle in sleep rested and sleep deprived young and older adults. Post-hoc analyses of the SLEEPY dataset revealed significant differences in LC fractional anisotropy between *sleep rested* and *sleep deprived* young adults, but not between *sleep rested* and *sleep deprived* older adults.

While fractional anisotropy tended to be higher in older than younger adults within the LC itself, older adults typically showed lower fractional anisotropy than younger adults along the noradrenergic bundle white-matter ascending tract, a typical age-related pattern in white matter (Medina & Gaviria, 2008; Sibilia et al., 2017; Sullivan & Pfefferbaum, 2006; Voineskos et al., 2012). The lack of associations observed in our datasets between LC fractional anisotropy and noradrenergic bundle fractional anisotropy may suggest these two regions are affected by diffusivity independently.

In the BASE-II and LEMON datasets, this age difference in the noradrenergic bundle contrasted with a lack of age differences in the right and left control white-matter frontopontine tracts, suggesting that the age effects in the noradrenergic ascending tract reflect more than just a global change in white matter. Thus, together, these data indicate that diffusivity properties of the LC and its ascending noradrenergic tract are affected by aging in opposite ways. Our findings of age differences in fractional anisotropy in the LC and its ascending tracts extend a growing set of observations of age differences in the structure of the LC in aging (Chen et al., 2014; Dahl et al., 2021; Dahl, Mather, Werkle-Bergner, et al., 2020; Langley et al., 2020; Morris, Tan, et al., 2020; Sun et al., 2019).

To date, most in vivo findings of LC structure have relied on LC-MRI sequences that show a cross-sectional increase in LC-neuromelanin sensitive contrast from young adulthood until around age 57, at which point it levels off or declines (Liu et al., 2019), potentially suggesting a gradual accumulation of neuromelanin followed by noradrenergic degeneration. One of the three data sets we examined (BASE-II) included neuromelanin-sensitive scans, and there were no significant correlations between LC-MRI contrast from those scans and diffusivity measures from the LC or noradrenergic bundle. This suggests that the diffusivity differences reflect different structural changes than the neuromelanin-sensitive scans. An important future research question is to expand the relationship between LC diffusivity measures and cognition, or markers of brain health, as has been found for LC-MRI contrast (Clewett et al., 2016; Dahl et al., 2019; Langley et al., 2020). One initial study along these lines found that medial and radial diffusivity in the LC-thalamus tract was correlated with memory performance in an older cohort (Langley et al., 2021).

Our results raise the question of what properties of the LC are changing to lead its tissue to show higher fractional anisotropy with age. One possibility could be an increase in inflammation that restricts fluid flow, as animal research has demonstrated that increases in microglial density affect diffusivity, as measured using an orientation dispersion index (Yi et al., 2019). In general, sleep disruption is associated with increased inflammation (Irwin et al., 2016). If greater inflammation within the LC were driving the baseline age differences seen in the BASE-II and LEMON data sets, we would expect sleep deprivation to make younger adults’ LC fractional anisotropy resemble that of older adults more. However, sleep deprived younger adults showed significantly lower LC fractional anisotropy than sleep rested younger adults.

Another possibility is that the age differences in LC diffusivity relate to age differences in LC tonic activity levels. Although still an open question, various findings suggest that the LC is more tonically active in aging (Mather, 2020; Weinshenker, 2018). Age differences in tonic levels of LC could contribute to differences in diffusivity as neuronal activity increases neuronal volume, while shrinking the volume of the surrounding fluid-filled spaces (Abe et al., 2017; Iwasa et al., 1980; Le Bihan et al., 2006; Nunes et al., 2021; Svoboda & Syková, 1991; Tirosh & Nevo, 2013) However, sleep reduces LC activity (Hayat et al., 2020; Khanday et al., 2016; Takahashi et al., 2010), so this potential mechanism would not be consistent with the opposing effects of aging and sleep deprivation.

Recent sleep deprivation studies looking at more global diffusivity have observed similar changes in fractional anisotropy (Elvsåshagen et al., 2015; Voldsbekk et al., 2021; Voldsbekk et al., 2020). Specifically, fractional anisotropy increased throughout the waking day, but decreased during sleep deprivation in relation to increased radial diffusivity and decreased axial diffusivity (Elvsåshagen et al., 2015; Voldsbekk et al., 2020). Notably, fractional anisotropy has been positively correlated with increased sleep quality (Khalsa et al., 2017).

More generally, older adults accrue sleep problems throughout life that eventually result in occasional to frequent bouts of sleep loss and the effects of sleep deprivation may not be as pronounced when compared to younger adults (Lavoie et al., 2018; Mander et al., 2016). For instance, adolescents experience greater changes in mood and anxiety after sleep deprivation than do older adults (Talbot et al., 2010). Likewise, young adults show more attentional deficits after sleep deprivation than do older adults (Zitting et al., 2018). The current results raise the question of whether some of the emotional changes seen after sleep deprivation in young adults are related to microstructural changes in the LC.

Mean and radial diffusivity in the LC also showed some age differences (results and tables provided in the supplementary material), although not as pronounced as fractional anisotropy. In the BASE-II dataset, mean diffusivity in the LC was significantly higher in younger adults, compared to older adults. In the LEMON dataset, mean diffusivity was significantly higher in the left LC of younger adults, compared to older adults. Though the cause for these laterality effects is not known, the BASE-II dataset is composed of mostly older adults, while the LEMON has more younger adults. Given the LC’s proximity to the fourth ventricle, older adults may be susceptible to neurodegeneration within the LC as well as partial volume effects (Langley et al., 2020; Liu et al., 2017; Sun et al., 2020).

Interestingly, with the SLEEPY Brain dataset, we observed expected age differences in the LC of *sleep rested* adults, with lower mean and radial diffusivities in younger adults, relative to older adults. In the *sleep deprived* condition, we did not observe significant differences in LC mean and radial diffusivities between age groups. These lack of LC differences in the age comparison of sleep deprived subjects is possibly due to *within* age group differences of *sleep deprived*, compared to *sleep rested,* young adults. Specifically, younger adults displayed higher mean and radial diffusivity in the LC when *sleep deprived*, compared with *sleep rested* young adults. Meanwhile, levels of diffusivity within older adults between *sleep rested* and *sleep deprived* conditions remained relatively unchanged.

Because the noradrenergic bundle overlaps with the LC atlas, we conducted an along-tract analysis for the noradrenergic bundle fractional anisotropy. As expected, we observed significantly lower FA in the first 10 points of the noradrenergic bundle, which anatomically approximately represent regions of the locus coeruleus, in younger adults compared with older adults. However, these significant differences were not observed as strongly in the SLEEPY dataset. We suspect this may be due to the low sample size in the SLEEPY dataset that is then further split by age and sleep condition. Changes in radial diffusivity along the noradrenergic bundle of cognitively impaired older adults from the Alzheimer’s Disease Neuroimaging Initiative were significantly greater, compared to healthy controls, around the area of the LC and again as the tract approached the hippocampus (Sun et al., 2020).

Early Alzheimer’s pathology associated with dysfunction of sleep has been suggested to appear decades before cognitive deficits (Benveniste et al., 2019; Cedernaes et al., 2017; Ehrenberg et al., 2018; Mander et al., 2016). Notably, in post-mortem brains, abnormal-tau pathology accumulates in the LC before appearing in cortical regions like the entorhinal cortex (Braak & Del Trecidi, 2015; Braak et al., 2011). Though we cannot make causal claims of our sleep related diffusivity findings, future studies could provide clinical insights with follow up studies. For example, by sleep depriving a large sample of young and older adults, specifically from REM sleep (most relevant to LC and emotional activity) and studying its effects on LC diffusivity and emotion, we could better understand the relationship between LC diffusivity and emotion regulation.

While most studies comparing diffusivity in younger and older adults focus on white matter, a growing number of studies have started to examine diffusivity differences in grey matter in cortical and subcortical nuclei. Patients with Alzheimer’s disease generally show less fractional anisotropy and greater mean diffusivity than age-matched healthy adults (Weston et al., 2015). However, studies following people with autosomal dominant familial Alzheimer’s disease have found increased mean diffusivity in grey matter regions during the pre-symptomatic period, and older adults with significant memory decline show lower diffusivity in the posterior cingulate/precuneus region (Jacobs et al., 2013). As Langley et al., suggested, age-related LC degeneration may result in restricted diffusion within older adults (Langley et al., 2020). Fractional anisotropy also shows a positive correlation with age in the caudate, putamen and globus pallidus in a healthy cohort aged 10-52 (Pal et al., 2011). Thus, the LC may not be the only brain region showing lower fractional anisotropy in older adults.

### 4.1 Limitations

The SLEEPY dataset is likely underpowered given the low sample size between sleep rested and sleep deprived groups and therefore all interpretations of our results should be considered for replication in a larger sleep-deprived dataset. However, this dataset is one of the few available sleep-diffusion public dataset, so these initial findings may provide motivation for future studies. Crossing fibers may also indicate opposite or unexpected relationships with diffusivity values that may be related to our unexpected findings (Lee et al., 2015; Oouchi et al., 2007). Despite the limitations of DTI, it remains a valuable tool that may help us to better understand the LC *in-vivo* within humans. In general, our datasets were comprised of younger and older adults that had no neurological or known sleep disorders and may not reflect the general aging population. We also did not examine axial diffusivity, which may be influenced by sleep deprivation (Elvsåshagen et al., 2015). Due to partial volume constraints, the locus coeruleus ROI may be contaminated by white matter and CSF (given the position near the 4^th^ ventricle). However, given the opposite findings in the ascending white matter tract, we were still able to extract meaningful signal.

## 5. Conclusions

In this study, we identified unique associations of LC diffusivity in the context of healthy adults across three different data sets. We consistently observed lower fractional anisotropy in the locus coeruleus of younger adults, compared to older adults but higher fractional anisotropy in the ascending noradrenergic bundle of younger adults, compared to older adults. Fractional anisotropy is a measurement of structural integrity, and these age findings add to a growing literature highlighting age-related differences involving the locus coeruleus. To our knowledge, this is the first study to compare diffusivity differences *in-vivo* in the locus coeruleus versus noradrenergic bundle. It also is the first to compare LC diffusivity differences between sleep rested and sleep deprived young and older adults.

## Supporting information

Supplemental Mean and Radial Diffusivity Results

## Disclosures and acknowledgments

No Disclosures. Research reported in this publication was supported by the National Institute on Aging of the National Institutes of Health under award number **R01AG025340**.

## Notes

### Competing Interest Statement

The authors have declared no competing interest.

