## Supplemental Mean and Radial Diffusivity Results for "Age Differences in Diffusivity in the Locus Coeruleus and its Ascending Noradrenergic Tract"

**Supplementary Results and Tables for Mean and Radial Diffusivities**

**1.1 Mean and Radial diffusivity show similar trends to fractional anisotropy.**

LC-MRI with mean or radial diffusivity correlation coefficients and 95% confidence intervals for younger and older adults are displayed in Tables 1 and 2, respectively. Like FA, we found no significant associations between LC-MRI and mean or radial diffusivity. Complete ANOVA tables for mean and radial diffusivity across datasets are displayed in Tables 3-8. We observed significant Age x ROI interactions in the BASE-II dataset for mean and radial diffusivities, $F(1.28, 380.26) = 38.34$, $p < .001$, $\hat{\eta}_G^2 = .05$, and $F(1.30, 387.03) = 44.29$, $p < .001$, $\hat{\eta}_G^2 = .057$, respectively (Table 3 and Table 6). We also observed significant Age x ROI x Hemisphere interactions for mean and radial diffusivities in the BASE-II dataset, $F(1.40, 414.51) = 24.32$, $p < .001$, $\hat{\eta}_G^2 = .15$, and $F(1.42, 420.66) = 23.81$, $p < .001$, $\hat{\eta}_G^2 = .016$, respectively.

Similarly, we observed significant Age x ROI interactions in the LEMON dataset for mean and radial diffusivities, $F(1.24, 267.66) = 34.65$, $p < .001$, $\hat{\eta}_G^2 = .064$, and $F(1.31, 282.94) = 41.16$, $p < .001$, $\hat{\eta}_G^2 = .073$, respectively (Table 4 and Table 7). We also observed significant Age x ROI x Hemisphere interactions for mean and radial diffusivities in the LEMON dataset, $F(1.41, 304.60) = 28.92$, $p < .001$, $\hat{\eta}_G^2 = .026$, and $F(1.46, 314.29) = 28.31$, $p < .001$, $\hat{\eta}_G^2 = .026$, respectively. Unlike the BASE-II dataset, we observed significant Age x Gender x ROI x Hemisphere interactions within LEMON mean diffusivity $F(1.41, 304.60) = 4.74$, $p = .019$, $\hat{\eta}_G^2 = .004$, and radial diffusivity, $F(1.46, 314.29) = 5.22$, $p = .012$, $\hat{\eta}_G^2 = .005$. For the SLEEPY Brain dataset, we observed a trending Age x ROI x Hemisphere x Sleep Condition interaction for mean and radial diffusivities, $F(1.25, 51.27) = 3.45$, $p = .06$, $\hat{\eta}_G^2 = .011$, and $F(1.28, 52.65) = 2.65$, $p = .101$, $\hat{\eta}_G^2 = .008$, respectively

(Table 5 and Table 8). Next, we report planned contrasts of mean and radial diffusivities across datasets.

Table 9 and Table 10 report BASE-II and LEMON means and 95% confidence intervals for mean and radial diffusivity within each ROI between young and older adults in each hemisphere. We observed varying results between BASE-II and LEMON diffusivities within ROIs. In BASE-II, mean diffusivity in the left locus coeruleus was not significantly different between young and older adults, but radial diffusivity was. BASE-II mean and radial diffusivities in the right locus coeruleus were significantly higher in younger adults, compared to older adults. In the left and right noradrenergic bundles, both mean and radial diffusivities were significantly lower in younger adults, compared to older adults. We observed no significant difference in mean and radial diffusivities between age groups in the BASE-II frontopontine tract.

In the LEMON dataset, we observed mean and radial diffusivity in the left locus coeruleus to be significantly higher in younger adults, compared to older adults. We did not observe any statistically significant differences between young or older adult mean and radial diffusivities in the right locus coeruleus. Younger adults, compared to older adults, had significantly lower mean and radial diffusivities in the left and right noradrenergic bundles. Except for significantly higher mean diffusivity in the right frontopontine tract of younger adults, compared to older adults, no other significant mean or radial differences were observed in the frontopontine tract between age groups.

Means and confidence intervals for the SLEEPY dataset mean and radial diffusivities are available in Tables 11 and 12.

In the *sleep rested* condition, we observed no significant differences in mean or radial diffusivities between young and older adults in the left locus coeruleus, $t(41) = -1.134$, $p = .263$, and $t(41) = -1.012$, $p = .317$, respectively. However, in the right locus coeruleus, younger adults had significantly lower mean and radial diffusivities, compared with older adults, $t(41) = -2.543$, $p = .014$, and $t(41) = -2.345$, $p = .023$, respectively. Both mean and radial diffusivities in the left noradrenergic bundle $t(41) = -2.283$, $p = .027$, and $t(41) = -2.122$, $p = .039$, respectively, and mean and radial diffusivities in the right noradrenergic bundle $t(41) = -3.185$, $p = .002$, and $t(41) = -3.417$, $p = .001$, were significantly lower in younger adults, compared to older adults. Similarly, mean, and radial diffusivities in the left frontopontine tract, $t(41) = -2.393$, $p = .021$, and $t(41) = -2.28$, $p = .027$, and right frontopontine tract, $t(41) = -2.595$, $p = .013$, and $t(41) = -2.593$, $p = .013$, were significantly lower in younger adults, compared to older adults.

Within the *sleep deprived* condition, we observed no significant differences in mean or radial diffusivities between young and older adults in the left locus coeruleus, $t(41) = 0.177$, $p = .860$, and $t(41) = 0.666$, $p = .509$, respectively. Nor did we observe significant differences between young and older adults' mean and radial diffusivities in the right locus coeruleus, $t(41) = 0.949$, $p = .348$, and $t(41) = 1.491$, $p = .143$, respectively. Both mean and radial diffusivities in the left noradrenergic bundle $t(41) = -2.371$, $p = .022$, and $t(41) = -2.728$, $p = .009$, respectively, and mean and radial diffusivities in the right noradrenergic bundle $t(41) = -2.859$, $p = .006$, and $t(41) = -3.265$, $p = .002$, were significantly lower in younger adults, compared to older adults. No significant differences in mean or radial diffusivities of the left frontopontine tract, $t(41) = -0.882$, $p = .387$, and $t(41) = -0.771$, $p = .444$, and right frontopontine tract, $t(41) = -1.679$, $p = .101$, and $t(41) = -1.588$ $p = .120$, were observed between younger and older adults.

**Table 1**

*Younger adults LC-MRI contrast correlations with confidence intervals*

| Variable | LC-MRI Contrast |
|---|---|
| Noradrenergic bundle MD – Left hemisphere | .04<br>[-.21, .29] |
| Noradrenergic bundle MD – Right hemisphere | .19<br>[-.06, .42] |
| Noradrenergic bundle RD – Left hemisphere | .10<br>[-.16, .34] |
| Noradrenergic bundle RD – Right hemisphere | .19<br>[-.06, .42] |
| Locus Coeruleus MD – Left hemisphere | -.01<br>[-.26, .24] |
| Locus Coeruleus MD – Right hemisphere | .05<br>[-.21, .29] |
| Locus Coeruleus RD – Left hemisphere | -.01<br>[-.26, .24] |
| Locus Coeruleus RD – Right hemisphere | .05<br>[-.20, .30] |
| Frontopontine MD – Left hemisphere | .11<br>[-.15, .35] |
| Frontopontine MD – Right hemisphere | -.02<br>[-.27, .23] |
| Frontopontine RD – Left hemisphere | .02<br>[-.23, .27] |
| Frontopontine RD – Right hemisphere | -.10<br>[-.34, .16] |

*Note.* Values in square brackets indicate the 95% confidence interval for each correlation.
The confidence interval is a plausible range of population correlations that could have caused the sample correlation (Cumming, 2014).

**Table 2**

*Older adults LC-MRI contrast correlations with confidence intervals*

| Variable | LC-MRI Contrast |
|---|---|
| Noradrenergic bundle MD – Left hemisphere | .06<br>[-.06, .19] |
| Noradrenergic bundle MD – Right hemisphere | .12<br>[-.00, .25] |
| Noradrenergic bundle RD – Left hemisphere | .06<br>[-.06, .19] |
| Noradrenergic bundle RD – Right hemisphere | .12<br>[-.00, .24] |
| Locus Coeruleus MD – Left hemisphere | -.02<br>[-.15, .11] |
| Locus Coeruleus MD – Right hemisphere | .10<br>[-.02, .23] |
| Locus Coeruleus RD – Left hemisphere | -.02<br>[-.15, .10] |
| Locus Coeruleus RD – Right hemisphere | .11<br>[-.02, .23] |
| Frontopontine MD – Left hemisphere | .02<br>[-.11, .14] |
| Frontopontine MD – Right hemisphere | .04<br>[-.08, .17] |
| Frontopontine RD – Left hemisphere | .06<br>[-.07, .18] |
| Frontopontine RD – Right hemisphere | .01<br>[-.11, .14] |

*Note.* Values in square brackets indicate the 95% confidence interval for each correlation.
The confidence interval is a plausible range of population correlations that could have caused the sample correlation (Cumming, 2014).

**Table 3**

*BASE-II Mean Diffusivity Mixed ANOVA*

| Variable | $F$ | $df_1^{GG}$ | $df_2^{GG}$ | $MSE$ | $p$ | $\hat{\eta}_G^2$ |
|---|---|---|---|---|---|---|
| Age (Younger, Older) | 7.11 | 1 | 297 | 0.00 | .008 | .007 |
| Gender (Female, Male) | 7.37 | 1 | 297 | 0.00 | .007 | .007 |
| ROI (LC, Noradrenergic Bundle, Frontopontine) | 1,376.75 | 1.28 | 380.26 | 0.00 | <.001 | .655 |
| Hemisphere (Left, Right) | 78.12 | 1 | 297 | 0.00 | <.001 | .025 |
| Age × Gender | 1.44 | 1 | 297 | 0.00 | .231 | .001 |
| Age × ROI | 38.34 | 1.28 | 380.26 | 0.00 | <.001 | .050 |
| Gender × ROI | 9.12 | 1.28 | 380.26 | 0.00 | .001 | .012 |
| Age × Hemisphere | 14.04 | 1 | 297 | 0.00 | <.001 | .005 |
| Gender × Hemisphere | 0.18 | 1 | 297 | 0.00 | .671 | .000 |
| ROI × Hemisphere | 217.13 | 1.40 | 414.51 | 0.00 | <.001 | .122 |
| Age × Gender × ROI | 0.02 | 1.28 | 380.26 | 0.00 | .939 | .000 |
| Age × Gender × Hemisphere | 1.02 | 1 | 297 | 0.00 | .313 | .000 |
| Age × ROI × Hemisphere | 24.32 | 1.40 | 414.51 | 0.00 | <.001 | .015 |
| Gender × ROI × Hemisphere | 0.22 | 1.40 | 414.51 | 0.00 | .722 | .000 |
| Age × Gender × ROI × Hemisphere | 0.14 | 1.40 | 414.51 | 0.00 | .789 | .000 |

**Table 4**

*LEMON Mean Diffusivity Mixed ANOVA*

| Variable | $F$ | $df_1^{GG}$ | $df_2^{GG}$ | $MSE$ | $p$ | $\hat{\eta}_G^2$ |
|---|---|---|---|---|---|---|
| Age (Younger, Older) | 6.18 | 1 | 216 | 0.00 | .014 | .008 |
| Gender (Female, Male) | 0.01 | 1 | 216 | 0.00 | .907 | .000 |
| ROI (LC, Noradrenergic Bundle, Frontopontine) | 2,354.77 | 1.24 | 267.66 | 0.00 | <.001 | .823 |
| Hemisphere (Left, Right) | 61.38 | 1 | 216 | 0.00 | <.001 | .026 |
| Age × Gender | 0.00 | 1 | 216 | 0.00 | .967 | .000 |
| Age × ROI | 34.65 | 1.24 | 267.66 | 0.00 | <.001 | .064 |
| Gender × ROI | 4.69 | 1.24 | 267.66 | 0.00 | .024 | .009 |
| Age × Hemisphere | 55.04 | 1 | 216 | 0.00 | <.001 | .023 |
| Gender × Hemisphere | 3.31 | 1 | 216 | 0.00 | .070 | .001 |
| ROI × Hemisphere | 314.66 | 1.41 | 304.60 | 0.00 | <.001 | .224 |
| Age × Gender × ROI | 1.10 | 1.24 | 267.66 | 0.00 | .309 | .002 |
| Age × Gender × Hemisphere | 3.66 | 1 | 216 | 0.00 | .057 | .002 |
| Age × ROI × Hemisphere | 28.92 | 1.41 | 304.60 | 0.00 | <.001 | .026 |
| Gender × ROI × Hemisphere | 1.33 | 1.41 | 304.60 | 0.00 | .260 | .001 |
| Age × Gender × ROI × Hemisphere | 4.74 | 1.41 | 304.60 | 0.00 | .019 | .004 |

**Table 5**

*SLEEPY Mean Diffusivity Mixed ANOVA*

| Variable | $F$ | $df_1^{GG}$ | $df_2^{GG}$ | $MSE$ | $p$ | $\hat{\eta}_G^2$ |
|---|---|---|---|---|---|---|
| Age (Younger, Older) | 11.07 | 1 | 41 | 0.00 | .002 | .074 |
| Sleep Condition (Rested, Deprived) | 1.45 | 1 | 41 | 0.00 | .235 | .010 |
| ROI (LC, Noradrenergic Bundle, Frontopontine) | 296.37 | 1.61 | 66.05 | 0.00 | <.001 | .787 |
| Hemisphere (Left, Right) | 10.93 | 1 | 41 | 0.00 | .002 | .017 |
| Age × Sleep Condition | 4.55 | 1 | 41 | 0.00 | .039 | .032 |
| Age × ROI | 0.43 | 1.61 | 66.05 | 0.00 | .610 | .005 |
| Sleep Condition × ROI | 5.02 | 1.61 | 66.05 | 0.00 | .014 | .059 |
| Age × Hemisphere | 2.89 | 1 | 41 | 0.00 | .097 | .005 |
| Sleep Condition × Hemisphere | 1.81 | 1 | 41 | 0.00 | .186 | .003 |
| ROI × Hemisphere | 21.67 | 1.25 | 51.27 | 0.00 | <.001 | .063 |
| Age × Sleep Condition × ROI | 1.98 | 1.61 | 66.05 | 0.00 | .154 | .024 |
| Age × Sleep Condition × Hemisphere | 2.41 | 1 | 41 | 0.00 | .129 | .004 |
| Age × ROI × Hemisphere | 0.22 | 1.25 | 51.27 | 0.00 | .699 | .001 |
| Sleep Condition × ROI × Hemisphere | 0.75 | 1.25 | 51.27 | 0.00 | .417 | .002 |
| Age × Sleep Condition × ROI × Hemisphere | 3.45 | 1.25 | 51.27 | 0.00 | .060 | .011 |

**Table 6**

*BASE-II Radial Diffusivity Mixed ANOVA*

| Variable | $F$ | $df_1^{GG}$ | $df_2^{GG}$ | $MSE$ | $p$ | $\hat{\eta}_G^2$ |
|---|---|---|---|---|---|---|
| Age (Younger, Older) | 3.52 | 1 | 297 | 0.00 | .062 | .003 |
| Gender (Female, Male) | 6.16 | 1 | 297 | 0.00 | .014 | .006 |
| ROI (LC, Noradrenergic Bundle, Frontopontine) | 992.17 | 1.30 | 387.03 | 0.00 | <.001 | .575 |
| Hemisphere (Left, Right) | 145.09 | 1 | 297 | 0.00 | <.001 | .047 |
| Age × Gender | 2.09 | 1 | 297 | 0.00 | .150 | .002 |
| Age × ROI | 44.29 | 1.30 | 387.03 | 0.00 | <.001 | .057 |
| Gender × ROI | 7.10 | 1.30 | 387.03 | 0.00 | .004 | .010 |
| Age × Hemisphere | 13.56 | 1 | 297 | 0.00 | <.001 | .005 |
| Gender × Hemisphere | 0.16 | 1 | 297 | 0.00 | .690 | .000 |
| ROI × Hemisphere | 184.32 | 1.42 | 420.66 | 0.00 | <.001 | .111 |
| Age × Gender × ROI | 0.09 | 1.30 | 387.03 | 0.00 | .829 | .000 |
| Age × Gender × Hemisphere | 0.79 | 1 | 297 | 0.00 | .376 | .000 |
| Age × ROI × Hemisphere | 23.81 | 1.42 | 420.66 | 0.00 | <.001 | .016 |
| Gender × ROI × Hemisphere | 0.28 | 1.42 | 420.66 | 0.00 | .677 | .000 |
| Age × Gender × ROI × Hemisphere | 0.26 | 1.42 | 420.66 | 0.00 | .695 | .000 |

**Table 7**

*LEMON Radial Diffusivity Mixed ANOVA*

| Variable | $F$ | $df_1^{GG}$ | $df_2^{GG}$ | $MSE$ | $p$ | $\hat{\eta}_G^2$ |
|---|---|---|---|---|---|---|
| Age (Younger, Older) | 4.62 | 1 | 216 | 0.00 | .033 | .006 |
| Gender (Female, Male) | 0.24 | 1 | 216 | 0.00 | .624 | .000 |
| ROI (LC, Noradrenergic Bundle, Frontopontine) | 1,499.72 | 1.31 | 282.94 | 0.00 | <.001 | .741 |
| Hemisphere (Left, Right) | 134.75 | 1 | 216 | 0.00 | <.001 | .060 |
| Age × Gender | 0.42 | 1 | 216 | 0.00 | .517 | .001 |
| Age × ROI | 41.16 | 1.31 | 282.94 | 0.00 | <.001 | .073 |
| Gender × ROI | 3.49 | 1.31 | 282.94 | 0.00 | .051 | .007 |
| Age × Hemisphere | 50.04 | 1 | 216 | 0.00 | <.001 | .023 |
| Gender × Hemisphere | 3.17 | 1 | 216 | 0.00 | .076 | .002 |
| ROI × Hemisphere | 329.55 | 1.46 | 314.29 | 0.00 | <.001 | .234 |
| Age × Gender × ROI | 1.00 | 1.31 | 282.94 | 0.00 | .338 | .002 |
| Age × Gender × Hemisphere | 4.58 | 1 | 216 | 0.00 | .034 | .002 |
| Age × ROI × Hemisphere | 28.31 | 1.46 | 314.29 | 0.00 | <.001 | .026 |
| Gender × ROI × Hemisphere | 1.41 | 1.46 | 314.29 | 0.00 | .244 | .001 |
| Age × Gender × ROI × Hemisphere | 5.22 | 1.46 | 314.29 | 0.00 | .012 | .005 |

**Table 8**

*SLEEPY Radial Diffusivity Mixed ANOVA*

| Variable | $F$ | $df_1^{GG}$ | $df_2^{GG}$ | $MSE$ | $p$ | $\hat{\eta}_G^2$ |
|---|---|---|---|---|---|---|
| Age (Younger, Older) | 8.78 | 1 | 41 | 0.00 | .005 | .061 |
| Sleep Condition (Rested, Deprived) | 1.48 | 1 | 41 | 0.00 | .231 | .011 |
| ROI (LC, Noradrenergic Bundle, Frontopontine) | 179.49 | 1.69 | 69.15 | 0.00 | <.001 | .685 |
| Hemisphere (Left, Right) | 13.46 | 1 | 41 | 0.00 | .001 | .024 |
| Age × Sleep Condition | 5.02 | 1 | 41 | 0.00 | .031 | .036 |
| Age × ROI | 1.11 | 1.69 | 69.15 | 0.00 | .328 | .013 |
| Sleep Condition × ROI | 5.54 | 1.69 | 69.15 | 0.00 | .009 | .063 |
| Age × Hemisphere | 2.22 | 1 | 41 | 0.00 | .144 | .004 |
| Sleep Condition × Hemisphere | 2.09 | 1 | 41 | 0.00 | .156 | .004 |
| ROI × Hemisphere | 23.83 | 1.28 | 52.65 | 0.00 | <.001 | .069 |
| Age × Sleep Condition × ROI | 2.58 | 1.69 | 69.15 | 0.00 | .092 | .030 |
| Age × Sleep Condition × Hemisphere | 1.70 | 1 | 41 | 0.00 | .200 | .003 |
| Age × ROI × Hemisphere | 0.22 | 1.28 | 52.65 | 0.00 | .704 | .001 |
| Sleep Condition × ROI × Hemisphere | 0.62 | 1.28 | 52.65 | 0.00 | .473 | .002 |
| Age × Sleep Condition × ROI × Hemisphere | 2.65 | 1.28 | 52.65 | 0.00 | .101 | .008 |

**Table 9**

*BASE-II Mean and Radial Diffusivity Means, Standard Error, Degrees of Freedom and 95% Confidence Intervals*

| BASE-II | Locus Coeruleus | | Noradrenergic Bundle | | Frontopontine Tract | |
|---|---|---|---|---|---|---|
| | Young Adult | Older Adult | Young Adult | Older Adult | Young Adult | Older Adult |
| Mean Diffusivity, Left Hemisphere | | | | | | |
| *M* [a] | 0.00148 | 0.00143 | 0.00079 | 0.00086 | 0.00071 | 0.00073 |
| *SE* | 3.00E-05 | 2.00E-05 | 1.00E-05 | 0 | 1.00E-05 | 0 |
| Lower CI | 0.00142 | 0.0014 | 0.00078 | 0.00085 | 7.00E-04 | 0.00072 |
| Upper CI | 0.00154 | 0.00146 | 8.00E-04 | 0.00086 | 0.00073 | 0.00073 |
| Mean Diffusivity, Right Hemisphere | | | | | | |
| *M* [a] | 0.00144 | 0.00136 | 0.00098 | 0.00126 | 0.00069 | 0.00069 |
| *SE* | 4.00E-05 | 2.00E-05 | 2.00E-05 | 1.00E-05 | 1.00E-05 | 0 |
| Lower CI | 0.00136 | 0.00132 | 0.00094 | 0.00124 | 0.00067 | 0.00068 |
| Upper CI | 0.00151 | 0.00139 | 0.00102 | 0.00128 | 0.00071 | 7.00E-04 |
| Radial Diffusivity, Left Hemisphere | | | | | | |
| *M* [a] | 0.00117 | 0.00109 | 6.00E-04 | 0.00066 | 0.00045 | 0.00046 |
| *SE* | 3.00E-05 | 2.00E-05 | 1.00E-05 | 0 | 1.00E-05 | 0 |
| Lower CI | 0.00111 | 0.00106 | 0.00059 | 0.00066 | 0.00044 | 0.00045 |
| Upper CI | 0.00124 | 0.00113 | 0.00061 | 0.00067 | 0.00047 | 0.00047 |
| Radial Diffusivity , Right Hemisphere | | | | | | |
| *M* [a] | 0.00114 | 0.00103 | 0.00081 | 0.0011 | 0.00048 | 0.00048 |
| *SE* | 4.00E-05 | 2.00E-05 | 2.00E-05 | 1.00E-05 | 1.00E-05 | 0 |
| Lower CI | 0.00107 | 0.00099 | 0.00077 | 0.00107 | 0.00046 | 0.00047 |
| Upper CI | 0.00122 | 0.00107 | 0.00085 | 0.00112 | 5.00E-04 | 0.00049 |

*Note.* CI = confidence interval.

[a] degrees of freedom = 297.

**Table 10**

*LEMON Mean and Radial Diffusivity Means, Standard Error, Degrees of Freedom and 95% Confidence Intervals*

| LEMON | Locus Coeruleus | | Noradrenergic Bundle | | Frontopontine Tract | |
|---|---|---|---|---|---|---|
| | Young Adult | Older Adult | Young Adult | Older Adult | Young Adult | Older Adult |
| Mean Diffusivity, Left Hemisphere | | | | | | |
| *M* [a] | 0.0015 | 0.00142 | 0.00076 | 0.00079 | 0.00073 | 0.00072 |
| *SE* | 2.00E-05 | 2.00E-05 | 0 | 0 | 1.00E-05 | 1.00E-05 |
| Lower CI | 0.00146 | 0.00138 | 0.00076 | 0.00078 | 0.00072 | 0.00071 |
| Upper CI | 0.00153 | 0.00147 | 0.00077 | 8.00E-04 | 0.00074 | 0.00073 |
| Mean Diffusivity, Right Hemisphere | | | | | | |
| *M* [a] | 0.00135 | 0.00135 | 9.00E-04 | 0.00113 | 0.00073 | 0.00071 |
| *SE* | 2.00E-05 | 2.00E-05 | 1.00E-05 | 1.00E-05 | 0 | 1.00E-05 |
| Lower CI | 0.00132 | 0.00131 | 0.00088 | 0.0011 | 0.00073 | 7.00E-04 |
| Upper CI | 0.00139 | 0.0014 | 0.00092 | 0.00115 | 0.00074 | 0.00072 |
| Radial Diffusivity, Left Hemisphere | | | | | | |
| *M* [a] | 0.00115 | 0.00106 | 0.00056 | 0.00059 | 0.00045 | 0.00045 |
| *SE* | 2.00E-05 | 2.00E-05 | 0 | 0 | 1.00E-05 | 1.00E-05 |
| Lower CI | 0.00112 | 0.00101 | 0.00056 | 0.00058 | 0.00043 | 0.00043 |
| Upper CI | 0.00118 | 0.0011 | 0.00057 | 6.00E-04 | 0.00046 | 0.00046 |
| Radial Diffusivity, Right Hemisphere | | | | | | |
| *M* [a] | 0.00101 | 0.00099 | 0.00073 | 0.00096 | 5.00E-04 | 0.00049 |
| *SE* | 2.00E-05 | 2.00E-05 | 1.00E-05 | 1.00E-05 | 1.00E-05 | 1.00E-05 |
| Lower CI | 0.00097 | 0.00094 | 0.00071 | 0.00094 | 0.00049 | 0.00047 |
| Upper CI | 0.00104 | 0.00103 | 0.00075 | 0.00099 | 0.00051 | 5.00E-04 |

*Note.* CI = confidence interval.

[a] degrees of freedom = 216.

**Table 11**

*SLEEPY Mean Diffusivity Means, Standard Error, Degrees of Freedom and 95% Confidence Intervals*

| SLEEPY | Locus Coeruleus | | Noradrenergic Bundle | | Frontopontine Tract | |
|---|---|---|---|---|---|---|
| | Young Adult | Older Adult | Young Adult | Older Adult | Young Adult | Older Adult |
| Left Hemisphere Rested | | | | | | |
| *M* [a] | 0.00165 | 0.00178 | 0.00083 | 9.00E-04 | 0.00083 | 0.00105 |
| *SE* | 1.00E-04 | 7.00E-05 | 2.00E-05 | 2.00E-05 | 7.00E-05 | 5.00E-05 |
| *Lower CI* | 0.00145 | 0.00164 | 0.00078 | 0.00086 | 0.00068 | 0.00094 |
| *Upper CI* | 0.00184 | 0.00192 | 0.00088 | 0.00093 | 0.00098 | 0.00115 |
| Left Hemisphere Deprived | | | | | | |
| *M* [a] | 0.00186 | 0.00184 | 0.00083 | 0.00089 | 0.00083 | 0.00091 |
| *SE* | 9.00E-05 | 7.00E-05 | 2.00E-05 | 2.00E-05 | 7.00E-05 | 5.00E-05 |
| *Lower CI* | 0.00167 | 0.0017 | 0.00078 | 0.00086 | 0.00069 | 0.00081 |
| *Upper CI* | 0.00204 | 0.00197 | 0.00087 | 0.00092 | 0.00097 | 0.00101 |
| Right Hemisphere Rested | | | | | | |
| *M* [a] | 0.00146 | 0.00183 | 0.00098 | 0.00114 | 8.00E-04 | 0.00101 |
| *SE* | 0.00012 | 8.00E-05 | 4.00E-05 | 3.00E-05 | 7.00E-05 | 5.00E-05 |
| *Lower CI* | 0.00122 | 0.00166 | 9.00E-04 | 0.00108 | 0.00066 | 0.00092 |
| *Upper CI* | 0.0017 | 0.002 | 0.00106 | 0.0012 | 0.00093 | 0.00111 |
| Right Hemisphere Deprived | | | | | | |
| *M* [a] | 0.00195 | 0.00182 | 0.00101 | 0.00114 | 0.00078 | 0.00091 |
| *SE* | 0.00011 | 8.00E-05 | 4.00E-05 | 3.00E-05 | 6.00E-05 | 5.00E-05 |
| *Lower CI* | 0.00172 | 0.00166 | 0.00093 | 0.00109 | 0.00066 | 0.00082 |
| *Upper CI* | 0.00217 | 0.00198 | 0.00108 | 0.0012 | 0.00091 | 0.00101 |

*Note. CI = confidence interval.*

[a] degrees of freedom = 41.

**Table 12**

*SLEEPY Radial Diffusivity Means, Standard Error, Degrees of Freedom and 95% Confidence Intervals*

| SLEEPY | Locus Coeruleus | | Noradrenergic Bundle | | Frontopontine Tract | |
|---|---|---|---|---|---|---|
| | Young Adult | Older Adult | Young Adult | Older Adult | Young Adult | Older Adult |
| Left Hemisphere Rested | | | | | | |
| *M* [a] | 0.00126 | 0.00139 | 0.00068 | 0.00075 | 0.00063 | 0.00085 |
| *SE* | 1.00E-04 | 7.00E-05 | 2.00E-05 | 2.00E-05 | 8.00E-05 | 5.00E-05 |
| *Lower CI* | 0.00106 | 0.00124 | 0.00063 | 0.00071 | 0.00048 | 0.00074 |
| *Upper CI* | 0.00147 | 0.00153 | 0.00073 | 0.00078 | 0.00079 | 0.00095 |
| Left Hemisphere Deprived | | | | | | |
| *M* [a] | 0.0015 | 0.00142 | 0.00067 | 0.00074 | 0.00063 | 7.00E-04 |
| *SE* | 9.00E-05 | 7.00E-05 | 2.00E-05 | 2.00E-05 | 7.00E-05 | 5.00E-05 |
| *Lower CI* | 0.00131 | 0.00129 | 0.00062 | 0.00071 | 0.00049 | 0.00059 |
| *Upper CI* | 0.00169 | 0.00156 | 0.00071 | 0.00077 | 0.00077 | 8.00E-04 |
| Right Hemisphere Rested | | | | | | |
| *M* [a] | 0.0011 | 0.00141 | 0.00084 | 0.001 | 6.00E-04 | 0.00082 |
| *SE* | 0.00011 | 8.00E-05 | 4.00E-05 | 3.00E-05 | 7.00E-05 | 5.00E-05 |
| *Lower CI* | 0.00088 | 0.00126 | 0.00077 | 0.00095 | 0.00046 | 0.00072 |
| *Upper CI* | 0.00132 | 0.00157 | 0.00092 | 0.00106 | 0.00074 | 0.00092 |
| Right Hemisphere Deprived | | | | | | |
| *M* [a] | 0.00159 | 0.0014 | 0.00087 | 0.00101 | 6.00E-04 | 0.00073 |
| *SE* | 1.00E-04 | 7.00E-05 | 4.00E-05 | 3.00E-05 | 7.00E-05 | 5.00E-05 |
| *Lower CI* | 0.00139 | 0.00126 | 0.00079 | 0.00096 | 0.00047 | 0.00063 |
| *Upper CI* | 0.00179 | 0.00155 | 0.00094 | 0.00106 | 0.00073 | 0.00082 |

*Note.* CI = confidence interval.

[a] degrees of freedom = 41.
